# Assembled chromosomes of the blood fluke *Schistosoma mansoni* provide insight into the evolution of its ZW sex-determination system

**DOI:** 10.1101/2021.08.13.456314

**Authors:** Sarah K Buddenborg, Alan Tracey, Duncan J Berger, Zhigang Lu, Stephen R Doyle, Beiyuan Fu, Fengtang Yang, Adam J Reid, Faye H Rodgers, Gabriel Rinaldi, Geetha Sankaranarayanan, Ulrike Böhme, Nancy Holroyd, Matthew Berriman

## Abstract

**Background:** *Schistosoma mansoni* is a flatworm that causes a neglected tropical disease affecting millions worldwide. Most flatworms are hermaphrodites but schistosomes have genotypically determined male (ZZ) and female (ZW) sexes. Sex is essential for pathology and transmission, however, the molecular determinants of sex remain unknown and is limited by poorly resolved sex chromosomes in previous genome assemblies.

**Results:** We assembled the 391.4 Mb *S. mansoni* genome into individual, single-scaffold chromosomes, including Z and W. Manual curation resulted in a vastly improved gene annotation, resolved gene and repeat arrays, trans-splicing, and almost all UTRs. The sex chromosomes each comprise pseudoautosomal regions and single sex-specific regions. The Z-specific region contains 932 genes, but on W all but 29 of these genes have been lost and the presence of five pseudogenes indicates that degeneration of W is ongoing. Synteny analysis reveals an ancient chromosomal fusion corresponding to the oldest part of Z, where only a single gene—encoding the large subunit of pre-mRNA splicing factor U2AF—has retained an intact copy on W. The sex-specific copies of U2AF have divergent N-termini and show sex-biased gene expression.

**Conclusion:** Our assembly with fully resolved chromosomes provides evidence of an evolutionary path taken to create the Z and W sex chromosomes of schistosomes. Sex-linked divergence of the single U2AF gene, which has been present in the sex-specific regions longer than any other extant gene with distinct male and female specific copies and expression, may have been a pivotal step in the evolution of gonorchorism and genotypic sex determination of schistosomes.

## BACKGROUND

*Schistosoma mansoni* is one of three main schistosome species that causes schistosomiasis, a neglected tropical disease that affects ∼240 million people worldwide [1]. Within the Phylum Platyhelminthes (flatworms), schistosomes are remarkable; while virtually all other flatworm families are hermaphrodites, family schistosomatidae are gonochoristic (separate sexes) and sexually dimorphic as adults. Sex is genetically determined with heterogametic females (2n=16, ZW) and homogametic males (2n=16, ZZ).

Adult female worms reside within the gynecophoric canal of adult males and the paired worms produce several hundred eggs a day. The eggs either traverse the intestinal wall to reach the lumen and be excreted in faeces or become trapped in host tissues, mainly liver and intestine, driving the pathology associated with schistosomiasis [2]. It has been postulated [3, 4] that dimorphism and gonochorism in schistosomes is an evolutionary adaptation to their residence in the venous system, close to capillary beds of warm-blooded host species; a division of labor between the sexes enables both a muscular male body to move against the blood flow of large veins and a thin slender female body shape to deposit eggs in small venules, allowing their efficient exit. However, the adaptions required to develop this dimorphism are unclear, limited by a lack of understanding of sex-linked molecular mechanisms, including unresolved sex chromosomes.

Despite major advances in the quality and quantity of published genome assemblies, sex chromosomes that are limited to the heterogametic sex (W and Y) are underrepresented in the growing list of whole genome assemblies. These sex-specific chromosomes are usually present at a lower copy number than autosomes, and the problem of assembling them is compounded by difficult to resolve highly repetitive sequences and by genetic divergence between the sex chromosomes, such that they can vary along their lengths [5]. There are exceptions—notably the recent publication of the eel genome [6] included resolved centromeres, subtelomeric sequences and the highly repetitiive Y chromosome short arm containing no gaps—but other sex chromosome assemblies, such as the *Drosophila* Y chromosome [7] and *Gallus gallus* W chromosome [8], are in fragmented states and even the reference human Y chromosome assembly [9] lacks continuity between the heterochromatic and euchromatic regions.

Degeneration of sex-limited chromosomes (W or Y) often distinguishes them from the shared (Z or X) chromosomes. Along the W chromosome of schistosomes, extensive heterochromatinization and the accumulation of satellite repeats, has been described, including a large satellite repeat SM-ɑlpha [10]. Extensive gene loss, or pseudogene-formation is also expected but without an adequate W assembly, it has not previously been possible to comprehensively describe the W-specific gene and repeat content that may play an important role in sex determination.

The *S. mansoni* genome was first published as a draft assembly [11], followed by a more contiguous version (v5) three years later [12] that took advantage of high throughput short-read sequencing technology. At that stage, as much as 80% of the genome had been assigned to chromosomes but gaps were prolific and large regions remained unresolved. The Z and W sequences were assembled together into merged scaffolds, with multiple Z-specific sequences and almost no resolution of W-specific sequences. As part of a sustained commitment to produce a complete genome sequence, in the present study, we have significantly improved upon previous efforts using a combination of long-read sequencing technology, optical mapping and manual curation to generate a highly contiguous chromosome-scale assembly that includes a fully assembled Z chromosome and a contiguous representation of the highly repetitive W chromosome. Our fully resolved reference genome is a key pre-requisite for understanding the evolution of sexual dimorphism in schistosomes and exposes sex-linked protein-coding and non-coding genes tentatively involved in sex determination.

## RESULTS

### The chromosome-level genome of *Schistosoma mansoni*

Using a combination of PacBio long-read and Illumina short-read sequencing, optical mapping, fluorescent *in situ* hybridization (FISH), Hi-C, and manual curation, we have assembled complete chromosomes from the 391.4 Mb genome of *S. mansoni*, including resolution of its Z and W sex chromosomes. The assemblies of chromosomes 2, 5, 6 and 7 comprise single scaffolds with telomeric repeats at either end; the remaining 5 chromosomes are also single scaffolds with a telomere at one end and sub-telomeric sequence at the other (Figure 1a,b).The number of gaps has decreased by 96% from 8,640 in the previous assembly to just 356 (Table 1).

**Figure 1:**
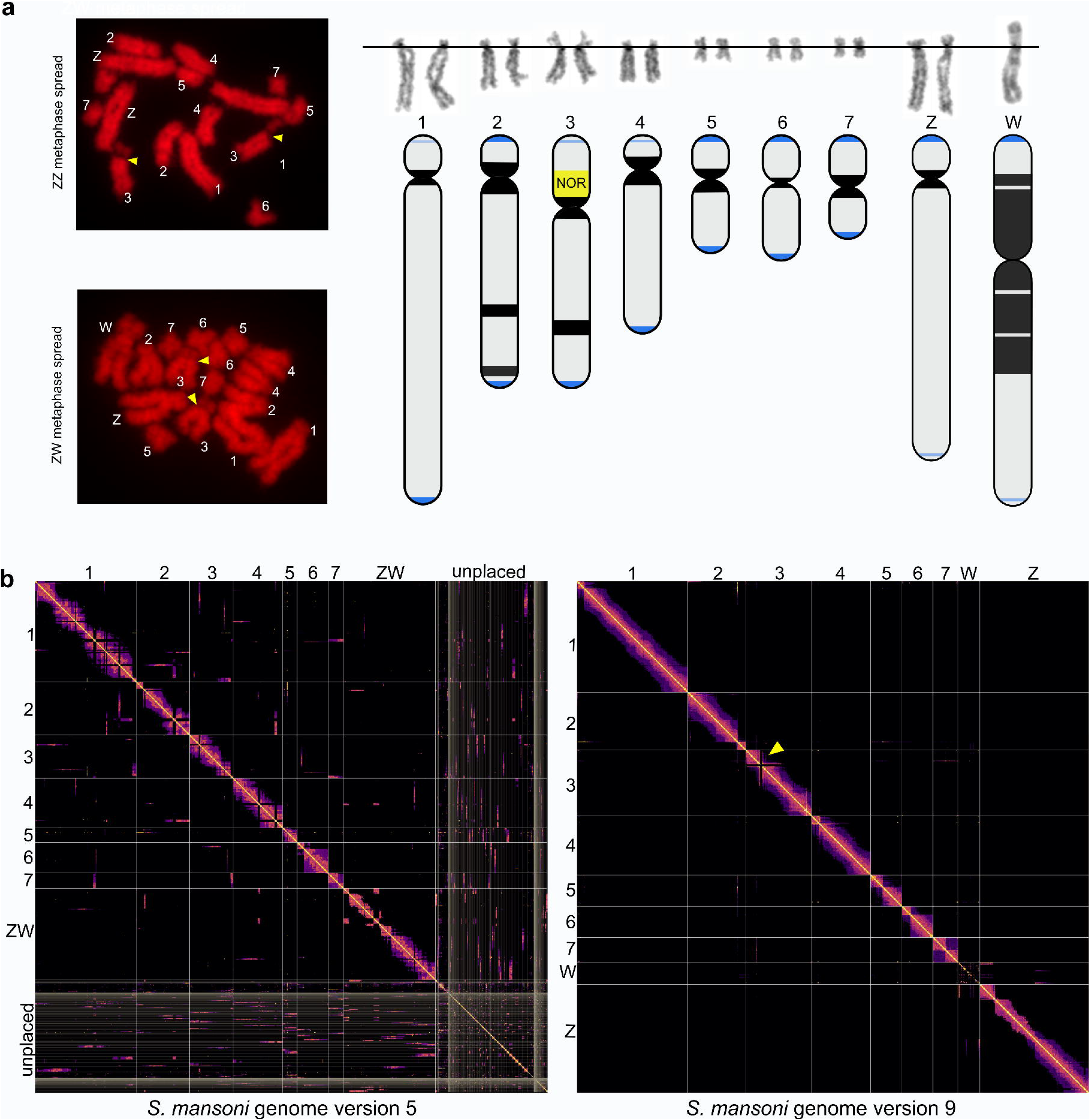
Ideograms of the *S. mansoni* chromosomes with HiC plots showing end-to-end chromosomal resolution. (a) Representative ZZ (male) and ZW (female) *S. mansoni* metaphase spreads, karyotypes, and ideograms. The yellow arrowheads point to the nucleolar organizer region (NOR; rDNA). Grey regions are euchromatic DNA, black are constitutive heterochromatin (C-band) regions, blue is confirmed telomeric sequence, and light blue bands are confirmed sub-telomeric sequence. (b) HiC visualization plots from genome version 5 (left) and version 9 (right) with the yellow arrowhead pointing to the NOR in chromosome 3.

**Table 1:** Genome-wide statistics for the S. mansoni haploid v9 assembly compared to the previous v5 assembly. The v9 assembly size has grown considerably with the addition of 26.9 Mb. The number of gaps present between versions was reduced by 96%, the majority of which are now only present in the collapsed repeat regions of the W chromosome. The chromosomes are assembled into 9 scaffolds (autosomes 1-7, W, and Z). Characterization of SLTS (spliced leader trans-splicing) in the transcripts has increased our previous estimates of only 7% of transcripts being trans-spliced to over 72% in the current assembly. *Completely new, previously partial, or previously unannotated.

|  | v5 | v9 |
| --- | --- | --- |
| Assembly size (Mb) | 364.5 | 391.4 |
| Gaps | 8,640 | 356 |
| Repeat Content (Mb) | 191.8 | 213.2 |
| **Scaffolds** |  |  |
| Number | 885 | 9 |
| N50 (Mb) | 32.1 | 52.8 |
| N90 (Mb) | 0.547 | 25 |
| Largest (Mb) | 65.5 | 89.1 |
| **Gene statistics** |  |  |
| Protein-coding genes | 10,116 | 9,794 |
| Novel genes* | - | 810 |
| Deleted genes | - | 867 |
| Pfam annotated | 66.9% | 70.3% |

| Transcript statistics |  |  |
| --- | --- | --- |
| Transcripts | 11,075 | 14,031 |
| Alternative splicing | 6.9% | 27.9% |
| Average exons per transcript | 5.9 | 7.9 |

The total repeat content of the assembly is 213.2 Mb (Table S1), a 21.4 Mb increase compared with the previously published version [12], reflecting the ability of PacBio long-read sequencing to account for repetitive regions that were previously difficult to assemble. For instance, an array of rRNA genes known as the nucleolar organizer region (NOR) of chromosome 3 (Figure 1) was highly collapsed in the earlier assembly and is now fully resolved. Newly resolved repetitive regions also include arrays of tandemly duplicated protein-coding genes enabling us to obtain a more accurate count for genes previously thought to be present as single copies. Two striking examples are the major egg antigens IPSE (IL-4-inducing principle of *S. mansoni* egg) and omega-1. These genes, specifically expressed in the eggs, have been intensely studied due to their roles in immune-modulation, pathogenesis and mechanisms of egg translocation to the intestinal lumen [13–16]. IPSE and omega-1 transcripts are encoded by paralogous gene arrays of at least 13 and 7 gene copies, respectively. In fact, based on the depth of coverage of aligned sequencing reads, these numbers are likely to be even higher and may contain as many as 20 and 14 copies of IPSE and omega-1, respectively (Figure S2).

We extended the analysis to identify other clusters of genes with conserved functions. Across the genome, there are 44 clusters of genes sharing similar predicted functions based on their protein (Pfam) domains, more than twice the number of clusters and domain types as seen in the previous v5 published genome version (Table S2). Clusters of *S. mansoni* Kunitz protease inhibitors and elastases are striking. Eleven Kunitz protease inhibitors (PF00014) exist in a cluster and 25 copies of elastase (PF00089; trypsin) are found across two clusters. The well-studied SmKI-1 (Smp_147730 in v5; Smp_311660, Smp_311670, and Smp_337730 in v9), is known to be involved in defense mechanisms of *S. mansoni* within the mammalian host [17]. The elastases are an expanded group of serine proteases originally noted for their role in host skin penetration, but are also expressed in intra-molluscan stages, where they likely facilitate movement of the parasite through snail tissue [18, 19].

### Annotation improvements through manual curation

We have significantly improved upon previous gene annotations of the *S. mansoni* genome. Using Augustus [20] and extensive RNA-seq evidence (Table S3) for gene prediction, followed by extensive manual curation, the total number of genes has decreased from 10,116 to 9,794 (excluding genes on scaffolds that correspond to alternative haplotypes; Table S4), compared to the v5 genome. This is the lowest number of genes for any sequenced platyhelminth; for instance, the cestodes *Echinococcus multilocularis* [21] and *Hymenolepis microstoma* [22] have 10,663 and 10,139 genes, respectively. In spite of the modest net reduction in genes, a total of 3,610 updates to gene models from v5 to v9 have been made, including 810 new, 867 deleted, 344 merged, 189 multiple copies, 190 split, and 1,210 with large structural changes (defined as >20% of coding region affected; Figure S3; Tables S5-S6). Using BUSCO v3.0.2 [23], the *S. mansoni* protein set was estimated to be 95.3% complete based on the representation of eukaryota orthologs (full genome-level BUSCO results at Table S7).

Spliced leader (SL) trans-splicing is an mRNA maturation process where an independently transcribed SL exon is transferred to a pre-mRNA. SL sequences originate from SL genes 613 bp in length, consisting of a 36 bp exon sequence (position 144–181 bp) flanked by an upstream precursor sequence (1–143 bp) and a downstream intron (182–613 bp) (Figure S1). A ∼1 Mb tandem array containing 41 full-length spliced-leader (SL) RNA genes has been resolved on chromosome 6 (Figure S1), together with an additional 109 partial gene sequences that contain the exon sequence only in the same array. On most other chromosomes, 1–4 SL gene fragments containing the exon sequence can also be found. Using RNA-seq data from all life cycle stages with an improved gene set (Table S3), we located SL receptor sequences in the transcripts of 6,641 genes in the primary assembly (i.e. no haplotypes), indicating that the majority of genes (66.3%) encode at least one trans-spliced isoform compared to 6.9% reported in the previous assembly (Table S8). This number is similar to the nematode *Caenorhabditis elegans* where ∼70% are identified as being trans-spliced [24].

The complexity of gene structures has increased substantially; the average number of exons per gene has increased from 5.9 to 7.9 (Table 1) and 97.7% of transcripts have both 5’ and 3’ untranslated regions (UTRs) annotated (Table S9). Further, the proportion of genes with alternative splicing to generate distinct transcribed isoforms has increased from 6.9% to 27.9%. Systematic improvements to gene finding and gene structural changes have enabled a richer set of putative functions to be ascribed to the *S. mansoni* proteome, reflected in the 47 new protein (Pfam) domains to *S. mansoni* from new genes and 79 new Pfams domains annotated in genes with improved gene structure (Table S10).

### Centromere motif conservation and divergence

*S. mansoni* chromosomes are monocentric [25], each with a cytologically distinguishable primary constriction (Figure 1a). The centromeric sequences are large repeat arrays that, on all chromosomes except 4 and Z, are highly conserved within a centromeric array and are between 93.1–98.5% similar to a 123 bp centromeric repeat proposed by Melters et al. [26] (Figure S4; Table S11). Between the centromeres of different chromosomes, the sequence conservation is more variable: 56% identity between the two most divergent centromere monomers (chromosomes Z and 6) and 100% identity between the centromeres of chromosomes 2 and 3 (Figure S4). The unit size is typical of the centromeric repeats of many other species [26], including the platyhelminth *Hymenolepis microstoma* [22]. The centromeric repeats for chromosomes 4 and Z have diverged from each other and from those of other chromosomes (Figure S4); their respective repeat units are 107 and 175 bp, and they are only 82 and 59% identical to the consensus from Melters et al. Centromeric repeats were previously estimated to comprise 0.48% of the genome (1.9 Mb) [26], but after including the divergent centromeres and estimating the degree to which all centromeric repeats were under-represented in the assembly based on mapped sequence coverage (from three PCR-free Illumina libraries), we estimate that centromeres make up at least 1.15% (4.5 Mb) of the genome.

### Architecture of the Z chromosome

Our new assembly includes a full-length 88 Mb Z chromosome that includes defined, recombining pseudoautosomal regions 1 (10.7 Mb) and 2 (42.9 Mb) and a non-recombining 33.1 Mb Z-specific region. In contrast to the previously published v5 assembly [12], where the Z chromosome was only partially resolved, the new sequence is 27.2 Mb larger with misassemblies corrected along its length, aided by the new long-range information that has been incorporated (Figure 2). In particular, the sequence that is unique to the Z chromosome (i.e., the Z-specific region, or ZSR), is clearly visible based on the lower depth of coverage of resequencing reads mapped from heterogametic females. The ZSR is flanked by two regions that are common to both sex chromosomes, termed pseudoautosomal region (PAR) 1 and 2.

**Figure 2:**
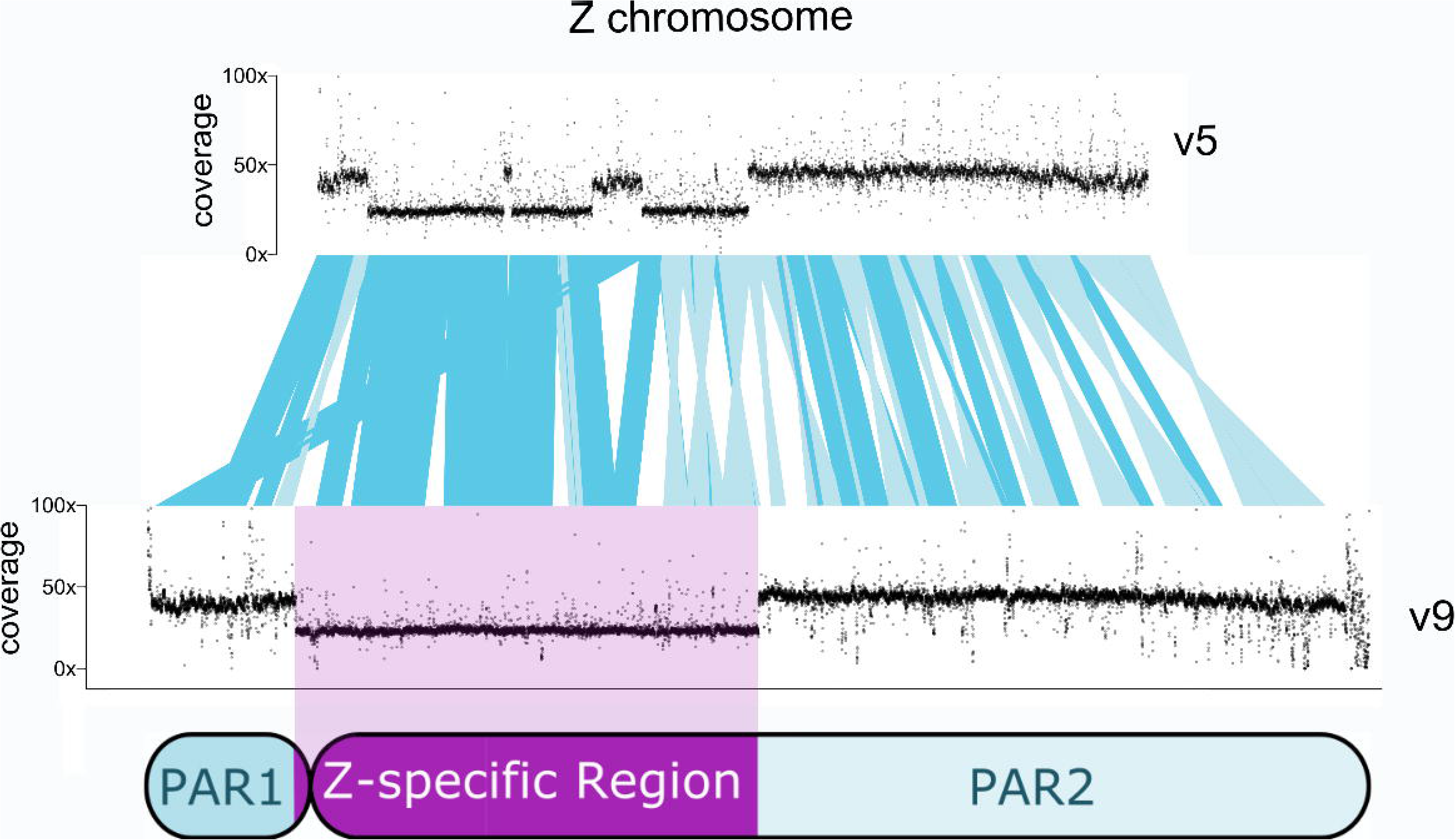
Improvements in the Z-specific region of the Z chromosome between the previous v5 *S.mansoni* genome assembly and current v9 assembly. Assemblies were compared using PROmer and visualized in ACT. The v5 assembly contained a partially resolved Z chromosome with misassemblies and between the Z-specific region (ZSR) and pseudoautosomal regions (PARs). Corrected inversions from v5 to v9 are shown in lighter blue. Coverage of mapped sequencing reads from female-only sequencing libraries highlight the ZSR as a region with approximately half the depth of coverage as pseudoautosomal regions.

Based on the earlier assembly (v5), it was previously shown [27] that the Z chromosome comprises different sub-regions or strata that have evolved differentially in the African and Asian Schistosoma lineages from a common ‘Ancestral’ stratum that is common to both lineages. Using the v9 assembly as a reference, where the ZSR is now resolved as a 33.2 Mb continuous sequence (Figure 2; Table S12), we plotted coverage of mapped sequencing reads across Z chromosome orthologs from four schistosome species (*S. mansoni*, *S. rodhaini, S. haematobium, S. japonicum*) and the hermaphroditic trematode *Echinostoma caproni.* In contrast to the relatively uniform mapped coverage for *E. caproni*, the ZSRs for the *Schistosoma* species are clearly visible, with a 19.1 Mb Ancestral shared region (ZSR2; ZSR coordinates 13,993,393-33,063,208) that has extended more recently in different directions amongst the African (*S. haematobium*, *S. rodhaini*, *S. mansoni)* and Asian species *(S. japonicum).* It also appears that in the Asian *S. japonicum*, two inversions have resulted in orthologues changing position and, therefore, creating coverage anomalies near the ZSR boundaries. The more recent 14 Mb African stratum (ZSR1; ZSR coordinates 1 - 13,993,392) extends beyond the centromere but is shorter than the Ancestral stratum (ZSR2). In contrast to the single, contiguous Z-specific region in the v9 assembly, the v5 assembly contained two blocks of what we now know to be PAR fragments which were incorrectly located inside the sex-specific region. It was previously reported that blocks of sequence shared by Z and W are located in the large region of recombination repression (i.e. the ZSR) [28]; based on this observation, Hirai, Hirai, and LoVerde [29] proposed three inversions in homologous Z/W regions from Z to W occurred before heterochromatinization, followed by at least one more inversion. These conclusions do not hold true in v9 and can now be attributed to misassemblies in v5.

To gain further insight into the evolutionary origins of the ZSR, we looked at the relationship between the Z chromosome and the chromosomal sequences of distantly related tapeworms. We have previously shown that flatworm genome structure can be defined based on conserved chromosome synteny blocks [30] (Figure 3b). When orthologs of *S. mansoni* and tapeworms are compared, synteny is largely preserved between these blocks, even though collinearity is disrupted. It is evident that one end of the Z chromosome is highly related to chromosome 3 of *Echinococcus multilocularis* and the other end is highly related to chromosome 5. When taken in isolation, the orthology evidence equally supports an ancient fusion in the schistosome lineage or an ancient fission in the tapeworm lineage. However, the position of the junction between the chromosome synteny blocks coincides with the position of the Ancient stratum (Figure 3a), suggesting that a fusion in the schistosome lineage is likely to have played a role, resulting in suppressed recombination.

**Figure 3:**
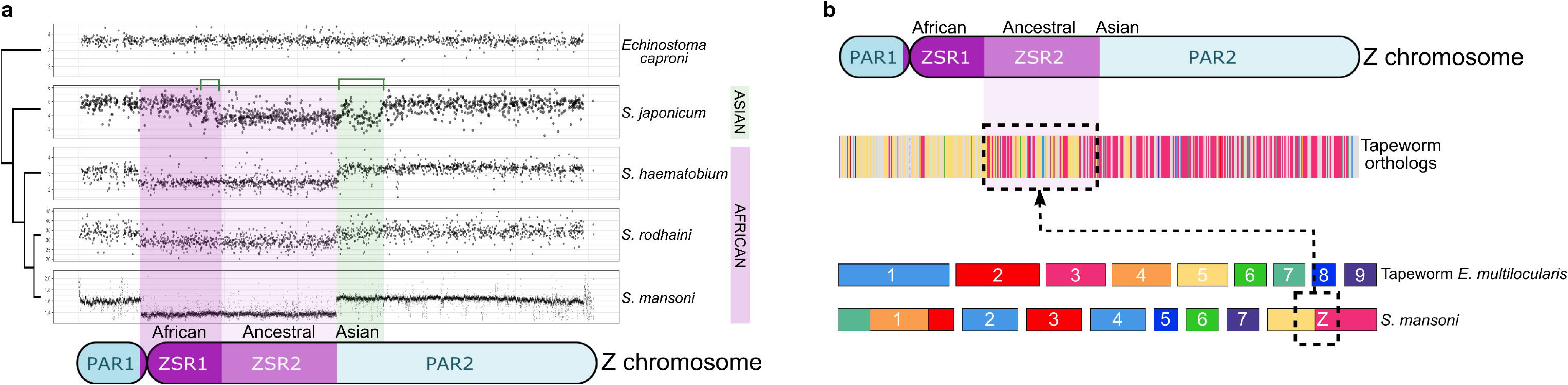
Z-specific regions of African and Asian *Schistosoma* spp. evolved differentially from an ancestral region that coincides with a likely fusion between chromosomes. (a) Evolutionary strata are defined through log_2_ genome coverage on the x-axis of one-to-one orthologs in four schistosomes and the hermaphroditic platyhelminth *Echinostoma caproni.* The African-specific stratum in dark purple defines Z-specific region 1 (ZSR1) of *S. mansoni* where approximately half coverage is seen in the African schistosomes *S. mansoni, S. rodhaini*, and *S. haematobium*. The Asian-specific stratum in green has reduced coverage specific only to *S. japonicum* with two possible inversions shown in green brackets. The ancestral *Schistosoma* stratum represents the schistosome orthologs ancestrally isolated to the Z sex chromosome between all schistosome species. (b) Tapeworm orthologs and chromosome synteny blocks show evidence of the fusion in the schistosome lineage between chromosomes 3 and 5 of the tapeworm *Echinococcus multilocularis.* Figure 3b adapted from Olson *et al* [30].

For neutral positions in the genome, the genetic diversity present is expected to reflect the number of copies of that region in the genome [31]. For the ZSR, the relative number of copies is 0.75 relative to autosomes (1.0), thus the diversity is expected to be lower than that of autosomes. Along the ZSR, we identified 352 genes in the African stratum and 580 in the Ancient stratum, which are flanked by 229 and 1,071 protein-coding genes in PAR1 and PAR2, respectively. We calculated the median nucleotide diversity (π) across the protein-coding genes of the autosomes and PARs and Z-specific regions (Figure 4; Table S13) using published genome variation data [32]. Across 50 kb windows, the autosomes have a median π range of 0.0026 to 0.0039. The PARs have a similar median π range to that of the autosomes at 0.0027 to 0.0032 in females and 0.0027 to 0.0034 in males suggesting that recombination between ZW and ZZ bivalents in the PARs is similar to that of the autosomal chromosomes. Also, the median π of the ZSR is significantly lower than that of the PARs for both males and females (p<0.001; Mann-Whitney test). We observed significantly lower π values in the Z African stratum when compared to the Z Ancestral stratum in both male and female samples (p<0.001; Mann-Whitney test), consistent with the effective population size of the Ancestral stratum being smaller for longer. The π values of the Z chromosome are close to that which would be expected in a neutral equilibrium with equal and constant male and female populations sizes (π_Z_/π_Autosomes_=0.75; [31] with π_Z_/π_Autosomes_=0.71 in males and π_Z_/π_Autosomes_=0.70 in females.

**Figure 4:**
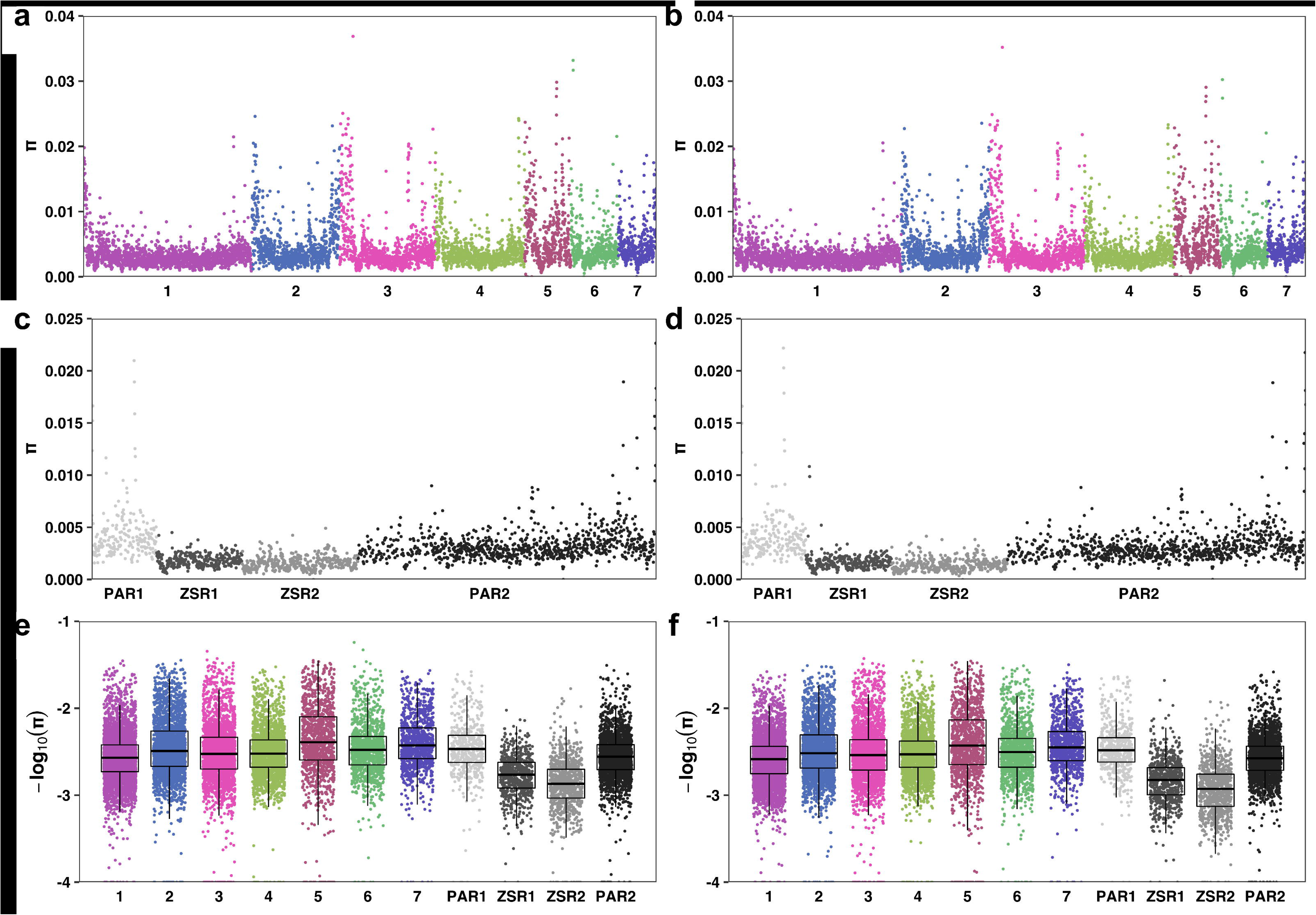
Median nucleotide diversity (π; pi) across the protein-coding genes of the autosomes, PARs, and Z-specific regions using published genome variation data. [32]. Median nucleotide diversity (π; pi) was calculated separately for males (left) and females (right) in 50 kb windows (a,b) or 5 kb windows (c-f) across all protein-coding genes. Pi is shown for the autosomes (a,b), PARs and ZSRs (c,d) and combined autosomal regions and ZSRs (e,f).

### Assembling the W Chromosome

The W chromosome shares >50 Mb of sequence with the Z chromosome in the pseudoautosomal regions, PAR1 and PAR2, that flank a highly repetitive W-specific region (WSR) (Figure 5; Table S12). In the v5 assembly, the highly repetitive W-specific region could not be resolved beyond ∼100 small and unordered contigs (1.1 Mb); by sequencing clonal females on multiple sequencing platforms, we resolved 22 repeat-rich W-specific scaffolds totalling 3.7 Mb (Figure S5). In many cases, the long reads used in our assembly were insufficient to fully span the arrays of repeats in the W chromosome. As a result, unique sequences are represented but the number of repeat units in many of the repeat arrays is vastly underestimated. After manual curation of the major repeat blocks, the W-specific assembly scaffolds were further ordered, oriented and linked by identifying as few as one, long PacBio subreads that spanned two consecutive blocks (Table S14). Metaphase FISH was also used to localize and orient three W-specific scaffolds that could not be placed through computational assembly methods (Figure S5).

**Figure 5:**
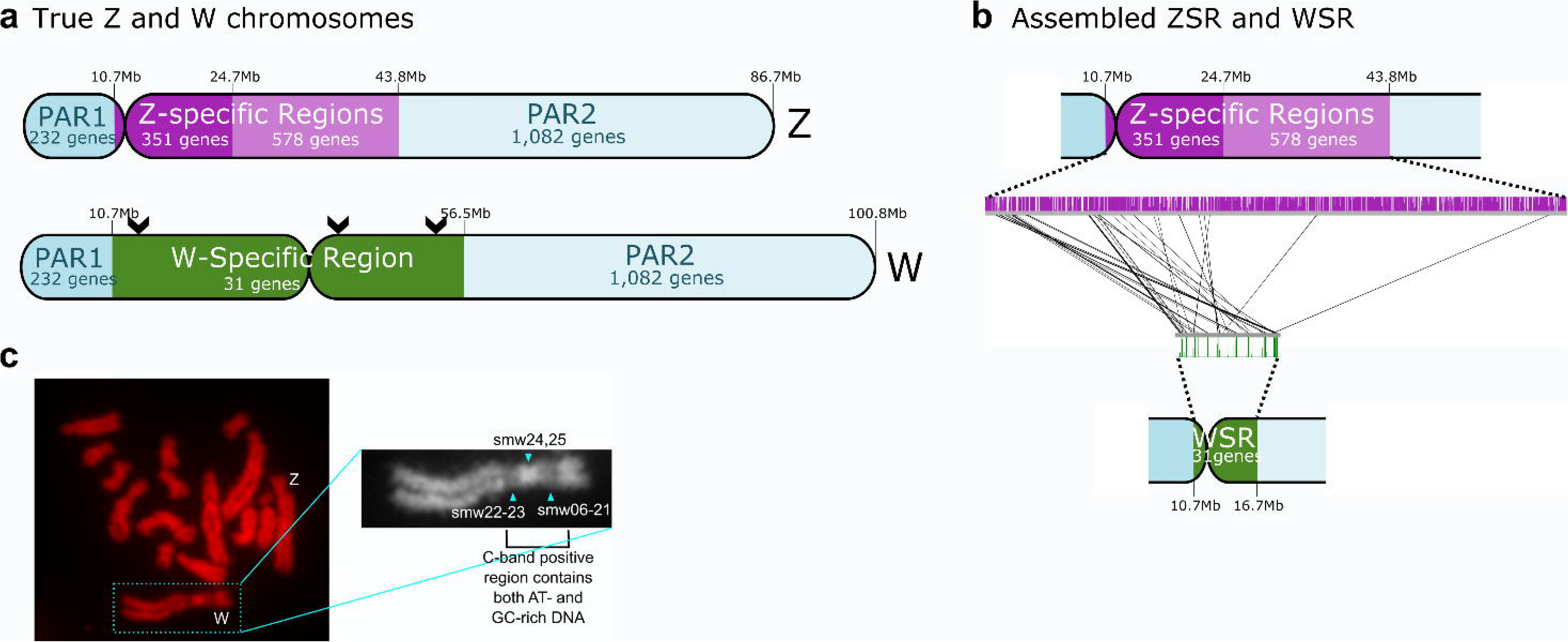
Detailed, annotated idiograms of the Z and W sex chromosomes. (a) The true size of the W chromosome is approximately 14% larger than Z which can be accounted for in the large expansion of repeats in the WSR. All but 36 genes have been lost on the WSR with 5 of those being pseudogenes and 2 present in triplicate and duplicate. Chevrons mark the approximate location of 3 euchromatin bands in the WSR. (b) The assembled size of the WSR is ∼6 Mb, less than its true size of ∼46 Mb because of 36 collapsed repeats. (c) C-banding shows the alternating AT- and GC-rich DNA repeats present in the WSR.

Previous karyotype measurements from 22 female metaphase cells [33] showed the W chromosome to be approximately 14% longer than the Z chromosome, a figure we confirmed with our own measurements of 14.7% using 6 female metaphase cells (Table S15; Figure S6). In particular, a long repetitive region in the short (p) arm of the W chromosome accounts for much of this size difference and is responsible for the p-arm being ∼40% of the W-chromosome length. Assuming a uniform density along the chromosome, relative measured lengths of other chromosomes with known assembly sizes (Figure S6), and genomic coverage of W-specific repeats (Table S16), we estimate the size of the W-specific region (WSR) to be ∼46 Mb. However, given that this region is heterochromatic and, therefore, more densely packed, its true size could be much longer. We attempted to estimate the degree to which repetitive regions remain collapsed within the assembly by mapping high-coverage Illumina sequencing reads from adult females. Extrapolating the read depth across repetitive regions (Table S16; see next section for results on W repeats) and comparing it with the median coverage for the genome (Table S17; ERS039722), we estimate a length of 17.6 Mb for the W-specific region. Clearly the mapping approach is inaccurate for estimating the true size of these collapsed regions. In fact, there are many regions of repetitive sequence in W where very few Illumina reads are mapped, indicating that certain repeat motifs are underrepresented in the sequence data. So-called “dark” and “camouflaged” regions of genomes have previously been reported, where specific sequencing technologies perform poorly (e.g. short tandem repeats, duplicated regions, regions with high GC content, non-random fragmentation) [34, 35].

### Repeat classification and heterochromatinization of the W chromosome

Like the human Y chromosome, the *S. mansoni* W chromosome is largely heterochromatic with a large proportion of its length composed of satellite repeats. There are just three bands of euchromatin on the W chromosome (chevrons in Figure 5) [10, 33]. Because some individual PacBio reads contained tandem arrays of the same repeat unit, we were able to assemble complete repeat units. Within the WSR constitutive heterochromatin, we characterized 36 unique repeats, named smw01-smw36 (Figure 5; Table S16). The 36 W-specific repeats comprise >95% of the assembled length of the W-specific region.

Of the 36 repeats, five (smw07, smw20, smw21, smw25, smw29) are related to the previously described 337 bp retrotransposable element SM*α*t-2 [36, 37]. Although a variant of SM*α*t-2 has been previously published as female-specific (SMAlphafem-1; NCBI accession U12442), we found one complete copy (coordinates: 23,37,004–23,936,670; 92.3% identity, 99.7% coverage, e-value 9.07e-133) and 38 partial copies (>75.0% identity; >95.0% coverage) on the Z chromosome. Metaphase FISH has shown striking fluorescence of a SM*α*t-2-related probe hybridizing near the short arm euchromatic gap [33, 37]. However, across the v9 genome, we found SM*α*t-2 repeats sporadically distributed on all autosomes and both sex chromosomes [38], but only as a large tandem array on the W chromosome, corresponding to the smw07 repeat found near the euchromatic band of the short arm [33].

Interestingly, 21 of the repeats can be grouped into five distinct families, where members within each family share at least 75.0% nucleotide identity, suggesting they may have evolved from a common ancestor including an SM*α*(aka SM-alpha and SMAlpha-fem) retrotransposon repeat family (smw03, 07, 20, 21, 29) (Table S16).

### Gametologues and their possible role in schistosome sex determination

The ZSR contains a total of 932 protein-coding genes. Of these, only 33 have clear homologous copies (termed gametologues) on the W chromosome, all within the WSR (Table S18). Although there is some positional clustering, extensive rearrangements by inversions, repeat expansions and transposable elements have largely disrupted collinearity between the WSR and the ZSR. The more recent African stratum contains 31 of the gametologues. For two of these, the corresponding W-copies have duplicated; there are three copies of genes encoding DnaJ domain proteins (heat shock protein 40 member B6) and two copies encoding a hypothetical protein with no discernible conserved features. At least five of the gametologues in the African stratum have degenerated into pseudogenes on W that have not yet been lost.

Considering the longest transcript for each gene, the W gametologues have an average of 55 amino acids less per protein sequence than the Z gametologues (Table S19). Only three W gametologues (spliceosome-associated protein, Smp_310950; ENTH domain-containing protein, Smp_303540; splicing factor U2AF 35 kDa small subunit, Smp_348830) are longer than their Z counterparts. Most Z and W gametologues are highly similar with average amino acid identities of >80% across their entire lengths using the Needle Wunsch algorithm in the EMBOSS package [39]. Excluding the five W pseudogenes and their Z gametologues, the gametologue pair with the greatest divergence was Smp_348820 on W and Smp_031310 on Z (encoding 40S ribosomal subunit S26) with only 47.6% identity. However, as with other low-similarity pairs, it was not possible, even through manual curation, to rule out gene finding inaccuracies due to a lack of isoform-specific transcript data.

We used previously published sex- and stage-specific RNA-seq [40, 41] to analyse differences in expression between the Z and W gametologue pairs (Figure 6). As expected, using unique mapping reads only for analysis, very few male reads mapped to the W gametologues. There were slight differences in the levels of expression between male and female samples for the Z gametologues, although RNA-seq coverage and replicate number from some of these samples were inadequate to enable robust analysis and interpretation. It has been shown one gametologue pair, encoding DnaJ homolog subfamily B member 4, have diel expression in males and females with the Z gametologue (Smp_336770) with the the Z gametologue cycling in adult females, males, and male heads, and the W gametologue (Smp_020920) cycling in females [42]. Expression of several W gametologues in female samples indicates possible stage-specific activity (such as Smp_317860, DnaJ heat shock protein family member B6) that is expressed in female larval cercariae and pre-dimorphic mammalian-stage schistosomula but not in adults; however, the Z gametologue to this gene, Smp_022330, shows consistent expression values across all stages.

**Figure 6:**
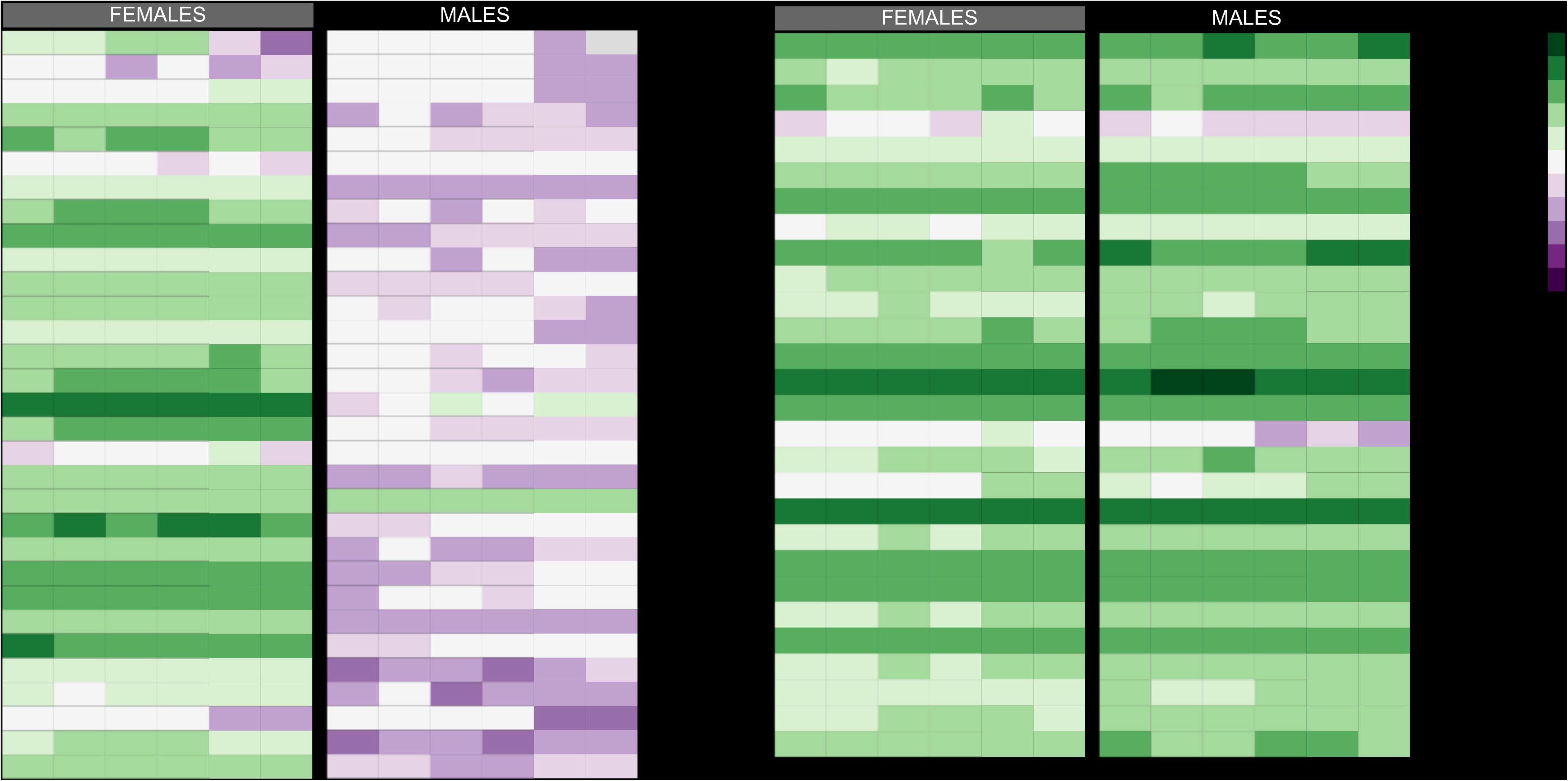
Illumina RNA-seq expression of the W and Z gametologues in adult paired and naive male and female *S. mansoni* worms. Unique mapping of RNA-seq to the gametologues reveals relatively similar expression of the Z gametologues between males and females for most gametologues. As expected, the W gametologues show expression limited to the female samples. Lines connect gametologue pairs between Z and W. In two cases, the W gametologue exists in triplicate or duplicate (see W gametologues Smp_317860, Smp_317870, Smp_348710 and Smp_318680, Smp_318860).

There is an almost complete lack of gametologues in the Ancestral stratum, which is consistent with this part of the chromosome having become sexually differentiated earlier and degenerative processes thus having been underway for longer. Within this long tract of degenerated sequence, two gametologues are clear exceptions. The first of these is a long multi-exon gene on Z, encoding a protein with ankyrin repeats and helicase domains. The corresponding gametologue on W is a pseudogene with several frameshifts and missing exons (Figure S7a). The second gametologue is predicted to encode the large subunit of splicing factor U2 snRNP auxiliary factor (Smp_019690 on Z and Smp_348790 on W). Strikingly, the sequences are almost identical (>95%) for most of their lengths but have divergent N-terminal sequences. After correcting for an artifactual frameshift in the W chromosome consensus sequence (based on aligned RNA-seq reads; Figure S7b), the copy on W shares the single-exon structure but the first 125 aa share only 45% identity.

## DISCUSSION

Our chromosome-scale assembly and curated annotation significantly extends the genetic resources for *S. mansoni*, and provides a more robust scaffold for genome-wide and functional genomic approaches for this important but neglected pathogen. It has enabled a greatly improved definition of the gene content, with the sequences of more than 25% of genes changed with >20% of coding region affected, and better resolution of those present in repetitive arrays, such as those encoding spliced leader RNA and stage-specific gene families. Amongst the gene families, many are known to encode highly abundant products —such as IPSE, omega-1, elastases, Kunitz protease inhibitors—that are important in host-parasite interactions. Major egg antigens omega-1 and IPSE are associated with a Th2 immune response in the host resulting in granulomatous inflammation around trapped parasite eggs [43]. Given the critical role of the intestinal granuloma for the egg translocation from the blood vessels to the intestinal lumen [44], genome expansions of these genes might have represented a selective advantage.

A major advance is in the analysis of schistosome sex chromosome evolution. Our previous analysis of orthologue synteny across the flatworms showed that the *S. mansoni* Z chromosome corresponds to two or more chromosomes in tapeworms [30]. From those data alone, it was not possible to determine whether a chromosome fusion had occurred in the schistosome lineage or whether it was a fission in tapeworms. However, in several other taxa, including filarial nematodes and several lepidotera, a chromosomal fusion has underpinned the genesis of sex chromosomes [45, 46]. We therefore speculate that a fusion has similarly occurred in the ancestral schistosome, creating a new pre-sex autosomal chromosome. The fusion event could have resulted in an isolated sex-determining locus that was advantageous to females and/or antagonistic to hermaphrodite worms. Consistent with this hypothesis, we show that the position of the putative fusion is within the oldest part of the Z-specific region of the chromosome and, within it, there is a single protein-coding ancestral gene (U2AF; splicing factor U2AF 65 kDa subunit) and a single pseudogene that are common to all African and Asian schistosomes. The alternative hypothesis to explain the observed synteny would require a fission at that position somewhere in the tapeworm lineage. This would have occurred prior to the formation of a sex determining region and the fission would, therefore, have played no role.

As one of two genomes found in the earliest-diverging part of the sex chromosomes, we identify the W gametologue encoding the pre-mRNA splicing factor U2AF 65 kDa subunit (Smp_348790) as a leading candidate gene for involvement in schistosome sex-determination. U2AF has been studied extensively in *Drosophila* for its association with the master sex-determining protein Sex-lethal (Sxl) [51] that is expressed exclusively in female flies. Sxl competes with U2AF binding to inhibit the splicing and translation of the *msl-2* gene (male-specific-lethal-2) [52, 53]. Considering that sex is determined by inhibiton of U2AF binding to pre-mRNA in *Drosophila*, it is tempting to speculate that the *S. mansoni* female-specific W copy of U2AF may antagonise the activity of the Z copy to inhibit the splicing of one or more genes. Further implicating U2AF in sex determination, the sex-specific regions also contain a homolog of the U2AF 35kDa subunit. In many taxa, U2AF is a heterodimer composed of large and small subunits that are required for spliceosome assembly in order to remove intron sequences from pre-mRNAs. U2AF binds to the 3’ splice site and polypyrimidine tract of introns in a complex with several other small nucleolar ribonucleoproteins (snRNPs) bound to the 5’ splice donor, commiting pre-mRNA to splicing (see review [50]). Our identification of U2AF2 is independently validated by Elkrewi et al. [49], who show using a search strategy based on the differential distribution of k-mers, that U2AF2 is the only intact gene in the ancient stratum of the ZSR.

How has sexual dimorphism evolved in schistosomatidae? The characterization of chromosomal fusions resulting the sex chromsomes, distinct evolutionary strata among closely related species, and the identification of U2AF allows us to propose a model of a model of the evolution of the schistosome sex chromosomes (Figure 8). At some point during the evolution of the Z and W sex chromosomes, the centromeric repeats diverged. It is not possible to know whether the centromere divergence occurred simply as a result of recombination or whether it played a more pivotal role in driving the suppression of recombination. Given the location of the centromere towards the far end of the more recent African stratum of the ZSR, the centromere divergence could have enabled a large expansion of the ZSR in the common ancestor of the African lineage of parasites. The high homology in amino acid sequence along with the conservation of functional domains between the gametologues suggests function has not changed between the gametologue pairs. Analysis of existing RNA-seq revealed sex- and stage-specific expression of the Z and W gametologues that could play a role in female-specific development. The duplication and triplication of two Z gametologues on W may be important in maintaining gene dosage or specialized female expression for those genes and is worthy of future study.

**Figure 7:**
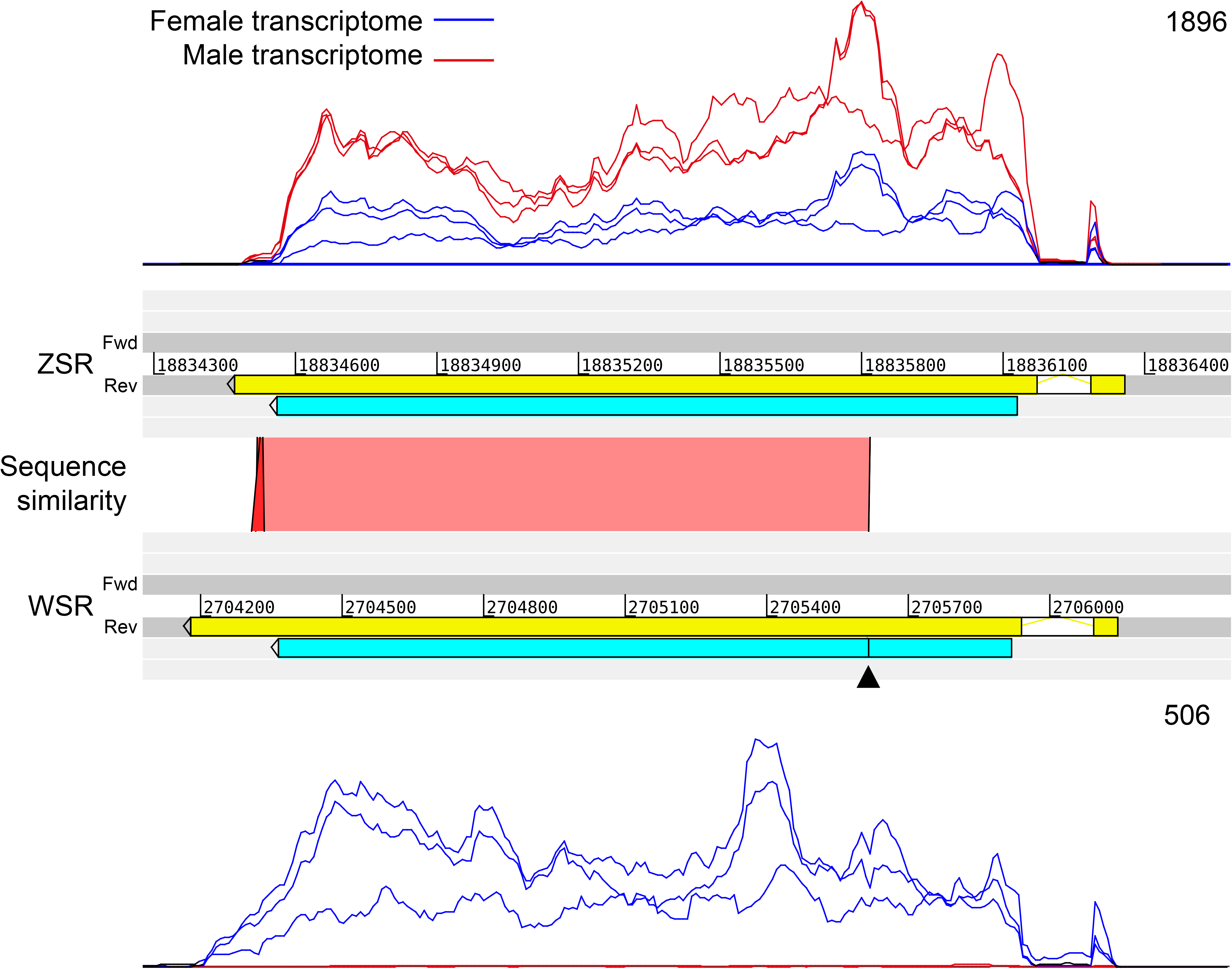
A comparison of U2AF 65 kDa subunit gametologues on the Z and W chromosomes. The gametologues of the large, 65 kDa subunit of U2AF (Z: Smp_019690; W: Smp_348790) are shown on ZSR (top) and WSR (bottom). Predicted transcript sequences in yellow. Sequence similarity was determined using PROmer and shows that the N-terminal region of the coding sequence (blue) is more diverse. The black arrow head highlights the position of a likely sequencing error on WSR which causes a frameshift, but which has been corrected in the gene model. Unnormalised coverage of RNA-seq reads is shown for female (bf_1, bf_2, bf_3) and male samples (bm_1, bm_2, bm_3). This highlights male expression on only the ZSR, with lower female coverage on ZSR and WSR as expected. Numbers above gene models indicate position on the contigs, numbers above RNA-seq coverage indicate maximum read depth.

**Figure 8:**
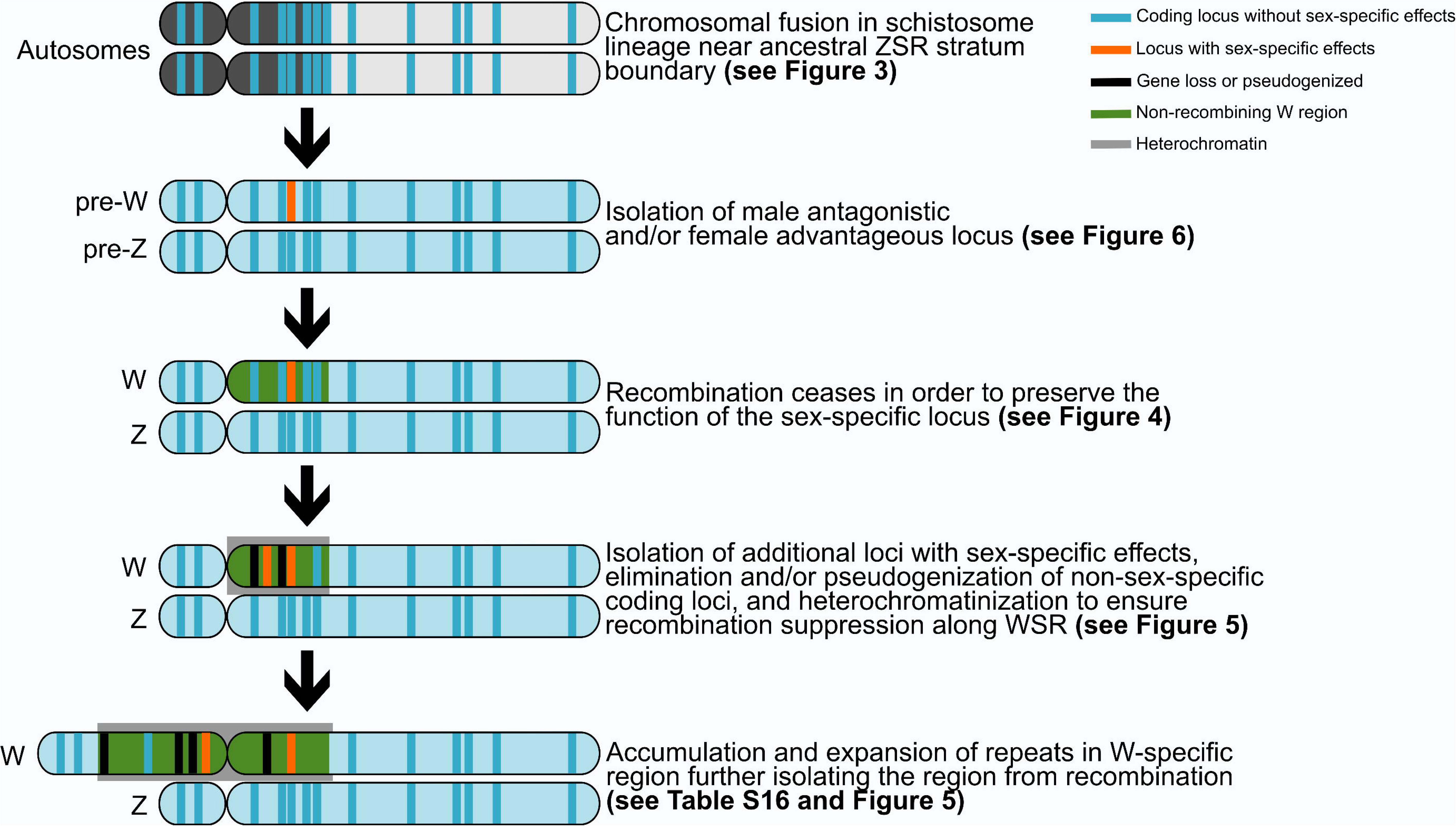
Hypothetical illustration of the schistosome Z and W sex chromosome evolution. A chromosomal fusion between two ancestral schistosome autosomes occurred near the ZSR stratum boundary (see Figure 3) creating a new set of autosomes. Followed by, or in conjunction with, this fusion event, a male antagonistic and/or female advantageous locus was isolated on the pre-sex chromsomes (see Figure 6; potentially pre-mRNA splicing factor U2AF). The need to isolate the phenotypic effects of the gene(s) in this locus on the pre-W chromosome required recombination suppression (see Figure 4). Isolation of additional loci with sex-specific effects and elimination and/or pseudogenization of non-sex-specific coding loci is evidenced in Fig 5. Following initial recombination suppression, extensive heterochromatinization of W ensured long-term recombination suppression between W and Z sex-specific regions and resulted in the huge expansion of repeats in the W-specific region (Figure 5; Table S16)

Although sexual dimorphism needs not rely on the existence of sex chromosomes and not all sexually dimorphic traits need to be linked to sex chromosomes [55], there must have been selective pressure to isolate sexually antagonistic and/or advantageous loci on non-recombining regions of sex chromosomes [56, 57]. Unlike many species in which a master sex-determining gene triggers male or female development, the absence of a W chromosome-specific genes suggests that multiple sex-determining loci were isolated on the sex chromosomes to produce separate sexes. With this in mind, we hypothesize that the W-copy of U2AF is regulating other gametologues or even genes located on the autosomes to control the suppression of male or female function. Identifying downstream interactions of U2AF with other genes is a critical next step for uncovering the mechanisms involved in schistosome sex determination. For example, do posttranslational modifications or splicing of W gametologues by U2AF directly inhibit the activity of a male-promoting product or create a male-lethal product? Future studies are needed to understand the functional role the gametologues like U2AF play in schistosome sex biology.

## CONCLUSIONS

*S. mansoni* is the most studied trematode and an accurate genome sequence underpins research into this important pathogen as well as enabling it to serve as a model for other trematodes. As the first species with completely assembled Z and W sex chromosomes, the *S. mansoni* genome provides a novel resource for studying other ZW organisms and is a crucial resource for future investigation into the sexual biology of schistosomes. The results presented provide a signfiicant advance toward understanding the evolution of sex chromosomes among the Schistosomatidae. As the agent of a prominent neglected tropical disease, understanding the evolutionary origins and molecular mechanism of sex determination in schistosomes may reveal new vulnerabilities to combat these parasites. The identification of the W-copy of U2AF as a candidate sex determining factor is clearly a major first step. This new assembly and annotation has already assisted in a broad range of studies on schistosomiasis including monitoring genetic diversity in field strains [32, 58], the discovery of alleles under selection for resistance to the antihelmintic praziquantel [59], and the analysis of stage- and sex-specific epigenetic changes [60–62]. Future studies using this resource will undoubtedly continue to reveal novel biological insights into schistosome development, infection, host-parasite interactions, and pathogenicity.

## METHODS

### Parasite material

#### *Schistosoma mansoni* developmental stages

A summary of the parasite material for genome and transcriptome sequencing can be found in Table S17 and Table S3, respectively. Unless otherwise specified, the different *S. mansoni* developmental stages were collected following described protocols [63, 64]. Unless otherwise noted, samples for RNA extraction were resuspended in 1 ml of TRIzol and stored at -80°C until a standard TRIzol RNA extraction method was performed. Genomic DNA was extracted using a standard phenol:chloroform DNA extraction method.

#### Sporocysts

Sporocysts were collected from Brazilian *B. glabrata* snails (BgBre) infected with 10 miracidia of their sympatric Brazilian *S. mansoni* (SmBre) strain. Secondary (daughter) sporocysts were dissected from 20 snails at 15 days and 4.5 weeks after infection. Following RNA extraction, DNA was removed with the Ambion® DNA-free™ Kit following the standard procedure and purified with the RNeasy® Mini Kit (QIAGEN).

#### Cercariae

At 4.5 weeks post exposure to 15-30 miracidia each, snails were washed, transferred to a beaker containing ∼50 ml conditioned water, and placed under light to induce cercarial shedding. Cercariae were collected and water was replaced every 30 minutes for 2 hours. Cercariae were incubated on ice for 30 minutes and concentrated by centrifugation at 1500 x g for 30 minutes at 4°C.

Snails exposed to single miracidium each were tested for patent infection after 5 weeks by exposure to light to collect genomic DNA from pooled male and pooled female cercariae. Snails with patent infection were kept and exposed to light every three days. Cercariae collected from each snail were stored for DNA extraction. Sex of the cercariae was identified by PCR [65].

#### Schistosomula and adult worms

Briefly, water containing cercariae was filtered, cercariae were washed, and tails were sheared off by ∼20 passes through a 22-G emulsifying needle. Schistosomula bodies were separated from the sheared tails by Percoll gradient centrifugation, washed, and cultured at 37°C under 5% CO_2_.

Adult worms were collected by portal perfusion from experimentally-infected mice at 6, 13, 17, 21, 28 and 35 days post infection following methods previously described [66]. Clonal female or male adult worms were collected from mice infected with PCR-confirmed female or male cercariae, respectively, shed from single monomiracium-infected snails.

For RNA preparation, samples were thawed on ice and transferred to MagNA Lyser Green Beads (Roche Molecular Systems, Inc). The samples were homogenized using the FastPrep-24 instrument (MB Biomedicals, UK) for two 20 second pulses with a speed setting of 6. A standard TRIzol RNA extraction followed and RNA was concentrated using RNA Clean and Concentrator Kit (Zymo Research) according to the manufacturer’s recommendations. RNA quality was assessed on the Bioanalyzer (Agilent) and samples with the highest quality were chosen for reverse transcription.

#### Miracidia

Livers were removed from hamsters 49 days post-infection with cercariae of the Liberian strain of *S. mansoni* and homogenised in PBS. The homogenate was centrifuged for 10 minutes at 5,500 x g at 4°C and the supernatant was discarded. The pellet was washed twice by resuspension in 0.9% NaCl followed by centrifugation as above. The pellet was resuspended in fresh conditioned water, exposed to light, and miracidia were collected. Miracidia were centrifuged for 30 minutes at 15,000 rpm at 4°C. Pelleted miracidia were resuspended in 100 µl TriFast (Peqlab) before storage at -80°C. The miracidia were allowed to thaw at room temperature before homogenisation with a polypropylene pestle, and snap frozen in liquid nitrogen. This was repeated twice more before TriFast was added to 500 µl. RNA was then extracted according to the manufacturer’s instructions. Extracted RNA was quantified using a BioPhotometer plus (Eppendorf). RNA quality was assessed with the Bioanalyzer RNA 600 Pico Kit (Agilent).

### Illumina and PacBio genome sequencing

Clonal male and female mate pair libraries (3 kb fragment size) were prepared from cercariae genomic DNA, following a modified SOLiD 5500 protocol adapted for Illumina sequencing [67]. Additionally, genomic DNA from clonal male and clonal female adult material was used to make separate PCR-free 400-550 bp Illumina libraries following previously described protocols [68], with the exception of using Agencourt AMPure XP beads for sample clean-up and size selection. Genomic DNA was precipitated onto beads after each enzymatic stage with an equal volume of 20% polyethylene glycol 6000 and 2.5 M sodium chloride solution. Beads were not separated from the sample throughout the process until after the adapter ligation stage. Fresh beads were then used for final size selection. Illumina libraries were sequenced on either a HiSeq 2000 or 2500 (Table S17).

Genomic DNA from *S. mansoni* clonal female adults was used to prepare a SMRTbell library following the Pacific Biosciences protocol ‘20 kb Template Preparation Using BluePippin Size-selection System’. The resulting library was used to produce 40 SMRT cells on the Pacific Biosciences RSII platform. We also prepared a PacBio library using genomic DNA from a pool of male cercariae from a snail monomiracidium-infection producing 28 SMRT cells on the Pacific Biosciences RSII platform (Table S17).

#### Optical mapping for genome assembly corrections and increased resolution

Female clonal cercariae were used to make agarose plugs using the CHEF Genomic DNA Plug Kit (Bio-Rad) following methods previously described [69]. High molecular weight *S. mansoni* genomic DNA was prepared by proteinase K lysis of trypsin-digested adults mixed with molten agarose set in plugs. DNA molecules were stretched and immobilized along microfluidic channels before digestion with the restriction endonucleases *BamHI* and *NheI*, yielding a set of ordered restriction fragments in the order that they occur within the genome.

The optical data was generated and analysed using the Argus Optical Mapping System from OpGen and associated MapManager and MapSolver software tools. As the *S. mansoni* genome is significantly larger than the 100 Mb cut-off suggested by OpGen for *de novo* assembly, OpGen’s GenomeBuilder software was used to generate targeted local optical map assemblies from the sequence contigs to provide additional mapping information. The median coverage of fluorescently-labelled molecules in the optical contigs from which consensus sequences were built was 30x. The raw data for each optical map contig were manually scrutinized using OpGen’s AssemblyViewer software, allowing us to validate accuracy (i.e. consistent coverage of ≥20x). Contigs with a visible dip in raw molecular coverage were discarded as assembly errors. This resulted in a set of manually curated, non-redundant optical contig consensus sequences that were generated near remaining scaffold gaps, rather than being generated to cover the whole genome, due to finite computational and analytical resources. Comparison of sequence contigs with validated optical contig consensus sequences allowed further scaffolding of the genome assembly and resolution of misassemblies as necessary in Gap5.

### *de novo* assemblies and manual curation

We combined existing short read data [11, 12] with additional Illumina data, long PacBio reads (Table S17), optical contigs, and genetic markers [70], to construct an intermediary genome assembly (version 7; GCA_000237925.3) that could be used by the public immediately while time-intensive manual curation took place. Misassemblies were corrected using long-read evidence, as well as optical map data and genetic markers [70]. Remaining gaps were filled using gap-filling software [71, 72]. Genetic markers [70] and an updated genetic linkage map(unpublished data, Chevalier et al) were used to assign further scaffolds to chromosomes, and to aid improvement and validation of the rest of the assembly. Version 7 contains 10 chromosomal scaffolds (8 chromosomes plus two scaffolds whose coordinates are known in the W chromosome; 95.91% of scaffolded bases), 13 scaffolds assigned to an autosome with known coordinates (11 of these are primarily repetitive scaffolds), 20 W-specific scaffolds without chromosomal coordinates, 17 scaffolds not assigned to a chromosome, and one mitochondrial scaffold.

Following the v7 assembly submission, we further improved the assembly, particularly in assembling all W-specific contigs and in creating individual chromosomal scaffolds for both Z and W sex chromosomes. To assemble the W chromosome, we first produced separate *de novo* assemblies for Illumina and then used Spades [73]) and CANU [74]) to assemble PacBio genomic reads that did not map to the v7 assembly with >500bp of soft-clipping. Second, the *de novo* assemblies were screened against the NCBI NR database in order to screen out any non- *S. mansoni* sequences. New contigs were examined in Gap5 [75] for absence of mapped reads from a male Illumina library (PCR-free pooled male cercariae) and presence of mapped reads from the PCR-free pooled female cercariae Illumina library (Table S17). Manual improvement was performed in Gap5 [75]. Putative new W-specific contigs were examined for sequence similarity to the 22 existing W-specific scaffolds in v7 to determine unique W-specific contigs. All genomic reads (Table S17) were re-mapped to the new assembly and concordant soft-clipped sequences were extended. This process was continued iteratively until no further progress could be made, by which point all contigs terminated in tandem repeats. At this point, the PacBio subreads were surveyed to find long read evidence linking the W chromosome tandem repeats together (Table S16). This elucidated the order of the repeats and W-specific regions to construct a single W chromosome scaffold.

Z and W-specific chromosomal regions were determined from mapping coverage of PCR-free female Illumina libraries (Table S17) with ∼22x coverage in the ZSR and ∼44x coverage in the PARs, as expected in ZW females. Female-only libraries were used to manually identify gametologues on the W chromosome.

We resolved the haplotypic diversity that typically exists in genome assemblies by sequencing clonal parasites derived from single miracidium-infected snails. Haplotype genes were determined in Gap5 [75] by identifying genes with half coverage, and localisation to a single scaffold that is also half coverage, as compared to non-haplotype scaffolds. An erroneously classified W chromosome scaffold (SM_V7_W019) from v7 was re-classified as a chromosome 1 haplotype. Haplotypes are represented in 259 scaffolds (2.74% of scaffolded bases) (Table S4; DOI:10.5281/zenodo.5149023).

#### Metaphase fluorescent *in situ* hybridization (FISH) to confirm order of W-specific scaffolds

*S. mansoni* NMRI strain daughter sporocysts from *B. glabrata* snails were dissected at 29 days post exposure. Sporocysts were placed in 0.05% (0.5mg/ml) colchicine (Sigma-Aldrich) and titurated ∼20 times using an 18G blunt-end needle. This single cell suspension was incubated at room temperature for 2-4 hrs to arrest cell division. Cells were spun at 500 x g for 5 min, incubated in nuclease-free water for 20 min at room temperature, and then preserved in ice-cold 3:1 methanol:acetic acid fixative.

Several primer sets were designed to amplify 15 kb-30 kb fragments using the 22 W-specific scaffolds identified post-v7. Fragments were amplified using either PrimeSTAR GXL polymerase (TaKaRa Bio) or LA Taq Hot Start Version Polymerase (TaKaRa Bio) per the manufacturer’s instructions. The PCR products were run on an agarose gel and bands of the targeted size were cut and isolated using the QIAEX II Gel Extraction Kit (Qiagen). We successfully amplified sufficient DNA for labelling for scaffolds W005, W002, and W014 to confirm their order in the v9 assembly (Figure S5). Multiplex metaphase FISH and karyotyping were done following the procedures previously described [76].

#### Arima-HiC data to validate the *S. mansoni* v9 assembly

The Arima-HiC Kit for Animal Tissues (Arima Genomics; Material Part Numbers: A510008 Document Part Number: A160140 v00 Release Date: November 2018) was used following the manufacturer’s instructions with ∼100 fresh female *S. mansoni* worms as input. An Illumina library was made using the Swift Biosciences Accel-NGSO 2S Plus DNA Library Kit, with the modified Arima Genomics protocol. The library was sequenced on the Illumina HiSeq X Ten platform resulting in high resolution with >260x coverage of the genome (Table S17). Arima-HiC data was aligned to the v7 assembly using BWA [77]; version 0.7.17). The HiC contact map was made with PretextMap (https://github.com/wtsi-hpag/PretextMap) and viewed in PretextView (https://github.com/wtsi-hpag/PretextView) (Figure 1). Minor misassemblies and placement of previously 31 unplaced scaffolds were done manually in Gap5 [75].

### Illumina RNA-seq and PacBio IsoSeq transcriptome sequencing across *S. mansoni* developmental stages

Illumina RNA-seq libraries were prepared with the TruSeq RNA Library Prep Kit following the manufacturer’s protocol. The Smart-seq2 protocol [78] was followed as described to synthesize full length cDNA from 1 µg total RNA for PacBio IsoSeq full-length transcript sequencing. cDNA was amplified in 12 cycles PCR and size fractionated in SageELF electrophoresis system (Sage Science). One or more cDNA size fractions were pooled for the library preparation. For some samples, libraries were produced from more of the size fractions obtained from the SageELF, with the aim of reducing size bias in the PacBio RSII sequencing reads (Table S3).

#### Heterozygosity in Z and W sex chromosomes and nucleotide diversity in the Z chromosome

Genome-wide SNP calling was performed using GATK HaplotypeCaller with PCR-free Illumina genomic libraries (Table S17) and 7 previously published samples (12663_1_4, 12663_2_4, 7164_6, 7164_7, 7307_7, 7307_8, 8040_3) [32].

To calculate nucleotide diversity (π), median and mean autosomal coverage was calculated for all samples in the Crellen *et al.* data set [32]. Individuals with >10x median and mean coverage on Z and W chromosomes were retained (54 male and 61 female). Of these, the ZSR:PAR ratio was calculated. Individuals with >0.70 ZSR:PAR ratio and a PAR/ZSR <1.5 were designated as males and individuals with <0.70 ZSR:PAR ratio and a PAR/ZSR >1.5 were designated as females. This resulted in a data set consisting of 54 males and 61 females. We used PIXY (v.0.95.01) [79] to calculate π in 50 kb sliding, non-overlapping windows across each chromosome separately for male and female populations for the autosomes. Nucleotide diversity for the ZSR and PARs was calculated in 5 kb sliding, non-overlapping windows. We then calculated the bootstrapped (95%) confidence intervals for each population median using 1000 bootstrap samples of genomic windows for each population using previously published methods [58] ( https://github.com/duncanberger/PZQ_POPGEN/blob/master/Figures/figure_2.md). We compared nucleotide diversity between ZSR and the PARs for male individuals testing for significance using an unpaired t-test.

#### W-repeat classification and quantification

Dot plots were generated for each repeat array on the W chromosome contigs to ensure that a representative repeat unit was selected from each visually distinct section of each repeat array. This process yielded 36 unique repeat unit sequences subsequently named smw01-smw36. The 36 repeat units were compared, pairwise, using blastn with a word size of 6 and dust off. For each comparison with an e-value <0.01, the percentage identity and bit score was recorded and plotted in a matrix plot to reveal similarities between repeat units that define repeat unit families (i.e. Sm-*α*).

An attempt was made to computationally quantify the W-repeats. Using female PCR-free Illumina data (sample 6520_5; Table S17), gDNA reads were mapped to 19 known single and multi-copy genes (e.g. SmVAL, omega-1) and to all 36 identified W-repeat sequences. Using bedtools coverage on 50 bp windows from the resulting bam file, the single-copy genes had a median coverage of 67 with a range of 54 to 72 and a median of median coverages of 67. SmVAL had double this (151x) and omega-1 had 10 times this (671x) as expected. Taking normal coverage to be 67x, W coverage should be half that at 33.5x. From this we calculated an estimated expected size for our W-repeats (Table S16).

### Gene finding

#### Protein-coding genes

A new protein-coding gene set was produced for the v9 assembly from evidence-based predictions from Augustus [20] with Illumina and PacBio transcriptome reads (Table S3), followed by manual curation. Repeat Modeller v2.0.1 [80] and Repeat Masker v4.1.2 on sensitive mode [81] were run to identify, classify, and mask repetitive elements, including low-complexity sequences and interspersed repeats. The masked genome was then used for gene finding with Augustus v3.2.2 [82] with the following parameters designed to predict one or more splice-forms per gene: *--species=schistosoma2 --UTR=1 --alternative-from-evidence=1.* To predict better gene models and alternative splicing, we used extrinsic information as evidence (i.e. ‘exonpart’ and ‘intron’ hints in Augustus) based on Illumina short reads of all life stages except egg (set priority = 4 in the hints file), and PacBio Iso-seq reads of three life stages (male, female and schistosomula; priority = 40) (Table S3).

To facilitate the comparison of gene sets between assemblies, we also transferred the latest gene models from v5 (based on GeneDB in July 2017) to v7 using RATT [83] with the PacBio setting. The transferred gene models were then compared to those from *de novo* predictions using gffcompare v0.9.9d [84], to determine consensus or novel transcripts (blastn hit of <94% coverage or nucleotide identity <78% between the two assembly versions). When changes occurred compared to a previous gene model, namely an amino acid sequence had changed >20% in either identity or coverage as determined by blastp, or the gene was merged with another gene, or split into several new genes, a new identifier (starting with Smp_3) was assigned and the old Smp number(s) was kept as a previous systematic id (PSID). Otherwise, the previous v5 Smp identifier was transferred to the v7 gene model. Genes that were related to retrotransposons in v5, or not transferred by RATT to the v7 assembly, were not kept in the new gene set. From v7 to v9, gene models were transferred using Liftoff [85]. For gene models with structural changes compared to the v5 gene set, or potentially novel genes predicted by Augustus in the v7 and v9 assemblies, we have carefully inspected them and curated them in Web Apollo [86] (Tables S5, S6).

For functional annotation, blastp v2.7.0 against SwissProt was used to predict product information, and InterproScan v5.25 [87]) to predict product protein domains and Gene Ontology terms. For some genes their product information was preserved from the v5 gene set (taken from GeneDB) if the evidence code was not “Inferred from Electronic Annotation”.

Coverage of UTRs in the genome sequence was calculated as following: first we extract the 5’- and 3’-UTR annotations from the gff file, adding up the total UTR length for each transcript, and then for each gene, we took the transcript with longest UTR as a representative. Finally, all UTRs were summed up for calculating the coverage. Other feature statistics were calculated using Eval v2.2.8 [88].

To recover possible additional novel genes from Boroni *et al* [89], the CDS/transcript sequences were obtained directly from the authors and aligned to the v9 gene set using blast, where genes with hits were considered as existing. For those without hits to current gene models, their sequences were aligned to the whole genome using blastn and PROmer [90]. Genes with hits to multiple scaffolds were discarded. For genes hitting to the same scaffold the overlapping hit regions were merged using “bedtools merge” and set as “exon” in a gff. All possible models were manually inspected in Apollo using the same RNA-seq tracks as in the publication. We found evidence for 8 of the 759 putative novel genes reported by Boroni *et al.* [89]) (Table S20).

We initially assessed genome completeness using BUSCO v3.0.2 [23]. Although only 85.8% complete eukaryota orthologs were found in the genome sequence (using “--mode genome”; Table S7), representation is expected to be considerably less than 100% in platyhelminths due to their phylogenetic distance from other species in the BUSCO databases [22]. It is known that BUSCO applied to genomic sequences underestimates the completeness of assemblies due to the difficulty of detecting complete genes in the assembly [91] providing further explanation for missing orthologs. As an alternative, we tested the completeness of our predicted gene models using BUSCO (“--mode proteins”) and recovered 95.3% complete eukaryota orthologs.

#### Transfer RNAs (tRNAs)

tRNAscan v.1.3.1 [92], was used to identify transfer RNAs (tRNAs) in the *S. mansoni* v9 assembly. The algorithm was run with default parameters except for “--forceow --cove”.

#### Long intergenic non-coding RNAs (lincRNAs)

In order to locate long intergenic non-coding RNAs in v9 of the *S. mansoni* genome assembly, we used RATT [83] to migrate previously generated annotation [93] from v5 to v9. To this end, we downloaded the published annotation as a GFF file, transformed it to EMBL file (as required by RATT) and proceeded to migrate the annotations using the “PacBio” setting of RATT. From a total of 7,029 lincRNAs annotated in v5, 6,876 transfers were made (6,874 unique, two duplications) and 273 lincRNAs were not transferred.

#### Spliced-leader RNAs (SL RNAs)

Using RNA-seq data (Table S3), we have located SL (spliced leader) sequences in 6,497 genes (Table S8) or 66.3% of all annotated genes in the primary assembly. SL sequences were identified using the canonical *S. mansoni* SL sequence AACCGTCACGGTTTTACTCTTGTGATTTGTTGCATG (Genbank M34074.1 [94]) and a custom in-house spliced leader detection script [95] (https://github.com/stephenrdoyle/hcontortus_genome/blob/5543173b7ee83b903d976931813d85f96f7a6e13/03_code/hcontortus_genome.section5_workbook.md). The script first trims a predefined SL sequence from the 5’ end of RNA-seq reads allowing for a minimum length match with an allowed error rate of 10% using Cutadapt [96]. The trimmed sequences are extracted, sorted, and counted, making a sequence logo. The trimmed reads are mapped to the genome using HiSat2 [97] and a BAM file of the mapped trimmed reads is generated for visualisation. A BED file is also made of the splice site coordinates along with a WebLogo [98] of 20 bp surrounding the splice site. Finally, the script determines the coverage of splice sites with transcript starts, (200 bp upstream and 30 bp downstream of the annotated start codon) and internal CDSs, accounting for both misannotated and internal splice variants.

Following published methods [30], we looked for alternative SL sequences using a custom python script to identify reads that (a) aligned to annotated genes, or within 500 bp upstream, and (b) were soft-clipped by more than 5 bp at the 5′ end relative to the annotated gene. Soft-clipped sequences were clustered using CD-HIT-EST v4.7 [99] and only one prominent cluster was identified. Thus, the *S. mansoni* SL sequence appears to be highly conserved within the genome, and there is only one sequence with the abundance of the known SL sequence, occurring in around 10% of the randomly chosen RNA reads.

#### Gene clusters and gene density in the *S. mansoni* genome

To explore whether there are particular gene functions overrepresented on some chromosomes, we searched for genomically adjacent genes (>=3) with the same Pfam annotations. To investigate whether gene families that had been incorrectly collapsed in the v5 assembly and are now expanded in the v9 assembly, this analysis was performed for both v5 and v9 using Pfam annotations from InterproScan (see “Protein coding genes” section above). For clusters with at least 5 genes, the start coordinates of the first and last genes as well as the number of genes were indicated (Table S2).

IPSE and omega-1 were found to be multi-copy genes clustered in two tandem repeat regions. In order to compare how many bases of curated IPSE and omega-1 genes could be mapped to the v5 and v7 assemblies, we ran Exonerate with a max intron size of 1,500 bp for both IPSE and omega. The IPSE gene Smp_112110 was used in Exonerate, but for omega-1, the mRNA sequence was used because the omega-1 gene has a long and complex gene structure. GFF files were produced of mapped features for IPSE and omega-1 which served to illustrate how many copies of these genes could be annotated.

The IPSE v9 sequence is 199,167 bp with the equivalent v5 sequence is 86,067 bp. The gap in v9 is approximately 29 kb larger than the total of the gaps in v5 in this region. There are approximately 84 additional kilobases in v9 in this region mostly due to expansion of repeat sequence to give a closer representation of reality (Figure S2). Likewise, the omega-1 v9 sequence is 155,103 bp and the v5 sequence is 105,726 bp. There is a 29,982 bp increase due to a large gap in v9, leaving 19,395 bp of additional sequence mainly due to expansion of the repeat array.

#### Gene expression across different *S. mansoni* life stages and sexes

To explore gene transcript levels across different life stages and between males and females, previously published RNA-seq data [40, 41] was used. Briefly, reads were mapped to *S. mansoni* v9 genome using STAR v2.4.2a [100]. Counts per gene and TPMs were summarised with StringTie v2.1.4 [101]. Mean TPM values were calculated for samples of the same life stage and sex and log-transformed. For gametologue expression, only unique mapping reads were used for quantification.

In comparing gene expression of gametologues on WSR and ZSR regions, the ACT genome browser [102] was used with PROmer version 3.07 [103] to show sequence similarity. A transposon inserted into the Smp_318710 pseudogenes was annotated based on PROmer sequence similarity to other transposons on ZSR. For Figure 7, bm_1, bm_2, bm_3 male libraries and bf_1, bf_2, bf_3 female libraries were used [104]. For Figure S7b, bf_1 was used.

### Identification of centromeres and telomeres

A 123 bp tandem repeat motif was identified in *S. mansoni* by Melters *et al* [26] due to its high abundance (∼1% of the genomic reads), relative to all other tandem repeats in the genome. The original consensus was derived from multiple chromosomes and an almost identical motif is present in chromosomes 1-3, 5-7, and W (Table S11). On both chromosome 4 and Z, single candidate tandem repeats were identified with broadly similar repeat lengths and sequences to previously described consensus motif [26].

We examined repeats in Gap5 [75], taking only the portion of the repeat with the centromere tandem repeat motif. Centromere size estimates (Table S11) were based on Illumina genomic sequencing from female clonally-derived cercariae (sample ERS039722 from Table S17) mapped to 1 representative repeat unit of each of the 8 centromere repeats. As a control, reads were also mapped to the 1st 121 bp of the genomic sequence covered by 12 known single copy genes. These 12 genes gave us a median coverage of 15x. From this we were able to extrapolate sizes for each of the 8 centromeric repeats which totalled 2.25 Mb.

A MAFFT/Jalview alignment was created from all centromere motif sequences [105] and a neighbor-joining tree was constructed using the ETE Toolkit Phylogenetic tree viewer [106] (Table S11). Centromere motif sequence similarity was assessed using the alignment tool PRSS with the Smith-Waterman algorithm (https://embnet.vital-it.ch/software/PRSS_form.html [107, 108].

Hirai and LoVerde [109] determined the sequence motif of schistosome telomeres (CCCTAA repeat) through FISH detection. In African schistosomes, the telomeric repeat sequence can be found in the heterochromatin and centromere of the W chromosome. Because it is theorized that *Schistosoma* originated in Asia (see review [110]), the African schistosomes experienced more gene shuffling than the Asian schistosomes, accounting for the presence of telomeric repeats outside the ends of the chromosomes [111].

## Abbreviations

PAR: pseudoautosomal region
WSR: W-specific region
ZSR: Z-specific region
BUSCO: Benchmarking Universal Single Copy Orthologs
Mb: megabase
Kb: kilobase
bp: base pair
aa: amino acid
tRNA: transfer RNA
lincRNA: long intergenic non-coding
SL: spliced leader
SLTS: spliced leader trans-splicing
NOR: nucleolar organizer region; rDNA

## Declarations

### Ethics approval

To propagate the life cycle of the *Schistosoma mansoni* NMRI strain (Puerto Rican) and obtain different developmental stages of the parasite, BALB/c mice and susceptible BB02 strain *Biomphalaria glabrata* snails are routinely infected with parasites at the Wellcome Sanger Institute (WSI). The mouse infections were conducted under Home Office Project Licence No. 80/2596 and No. P77E8A062, and all protocols were presented and approved by the Animal Welfare and Ethical Review Body (AWERB) of the WSI. The AWERB is constituted as required by the UK Animals (Scientific Procedures) Act 1986 Amendment Regulations 2012. With the exception of sporocysts and miracidia, all life cycle stages were collected at the WSI.

*Schistosoma mansoni* SmBRE strain sporocysts dissected from infected BB02 *B. glabrata* snails were collected at The University of Perpignan laboratory which has permission A 66040 from both the French Ministère de l’agriculture et de la pêche and the French Ministère de l’Education Nationale de la Recherche et de la Technologie for experiments on animals and certificate for animal experimentation (authorization 007083, decree 87-848 and 2012201-0008) for the experimenters. Housing, breeding and animal care follow the national ethical requirements.

*Schistosoma mansoni* NMRI strain miracidia were collected at Justus-Liebig-University Giessen Institute for Parasitology. Animal experiments were approved by the Regional Council (Regierungspräsidium), Giessen, Germany (V54-19 c 20/15 c GI 18/10), which are in accordance with the European Convention for the Protection of Vertebrate Animals used for experimental and other scientific purposes (ETS No 123; revised Appendix A).

### Consent for publication

Not applicable

### Availability of data and materials

The primary genome assembly generated and analyzed during this study are available on the European Nucleotide Archive (ENA) website under project accession PRJEB13987. Additional assembled haplotypes, haplotype annotations, and primary assembly annotations can be found in permanent links at WormBase ParaSite under BioProject PRJEA36577 and at Zenodo DOI:10.5281/zenodo.5149023. All other data generated or analyzed during this study are included in this published article and its supplementary information files.

## Competing interests

The authors declare that they have no competing interests.

## Funding

This work was supported by the Wellcome Trust grants 098051 and 206194. For the purpose of Open Access, the authors have applied a CC BY public copyright licence to any Author Accepted Manuscript version arising from this submission.

## Authors’ contributions

MB designed research, which was coordinated by NH; SKB and GR maintained the parasite life cycle and generated parasite material, SKB, DB, and GS prepared genomic DNA, RNA and transcriptome sequencing libraries; BF, FY, and SKB performed FISH experiments; SKB, AT, ZL, SD, DB, FR, AJR, UB analyzed data; SKB drafted the complete manuscript, with sections of text contributed by AT, AJR and ZL. All authors read and approved the final manuscript.

## Acknowledgements

The *Schistosoma mansoni* genome project is funded by the Wellcome Trust through their core support of the Wellcome Trust Sanger Institute (grant 098051 and 206194). We thank Simon Claire and his team for technical assistance with the life cycle maintenance and collection of parasite material. We thank Tom Huckvale, Hayley Bennet, and Arporn Wangwiwatsin for technical support in generating RNA-seq and gDNA libraries. Thank you to Christoph Puethe for assisting in the ENA submission. A special thank you to Karl Hoffman and Josephine Ford-Thomas for supplying additional infected snails used for metaphase FISH, Paul Brindley and Victoria Mann for clonal worms, and Christoph Grunau and Benjamin Gourbal for providing *in vivo* sporocysts for RNA-seq.

## Additional Files

Additional file 1: **Supplementary Tables S1 to S20**

### Supplementary Figures Titles and Legends

**Figure S1:**
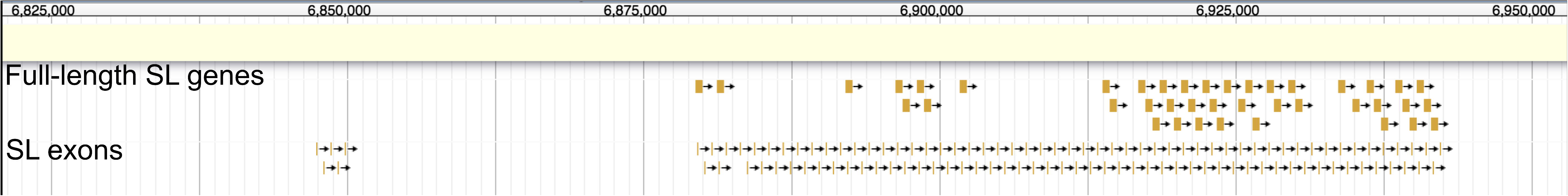
Array of spliced-leader RNA genes on chromosome 6 of the *S. mansoni* genome. On chromosome six, a 62.6 kb locus exists containing 41 full-length spliced leader RNA genes (top track). An additional 109 partial gene sequences that contain the spliced leader exon sequence only exist in the same array (bottom track).

**Figure S2:**
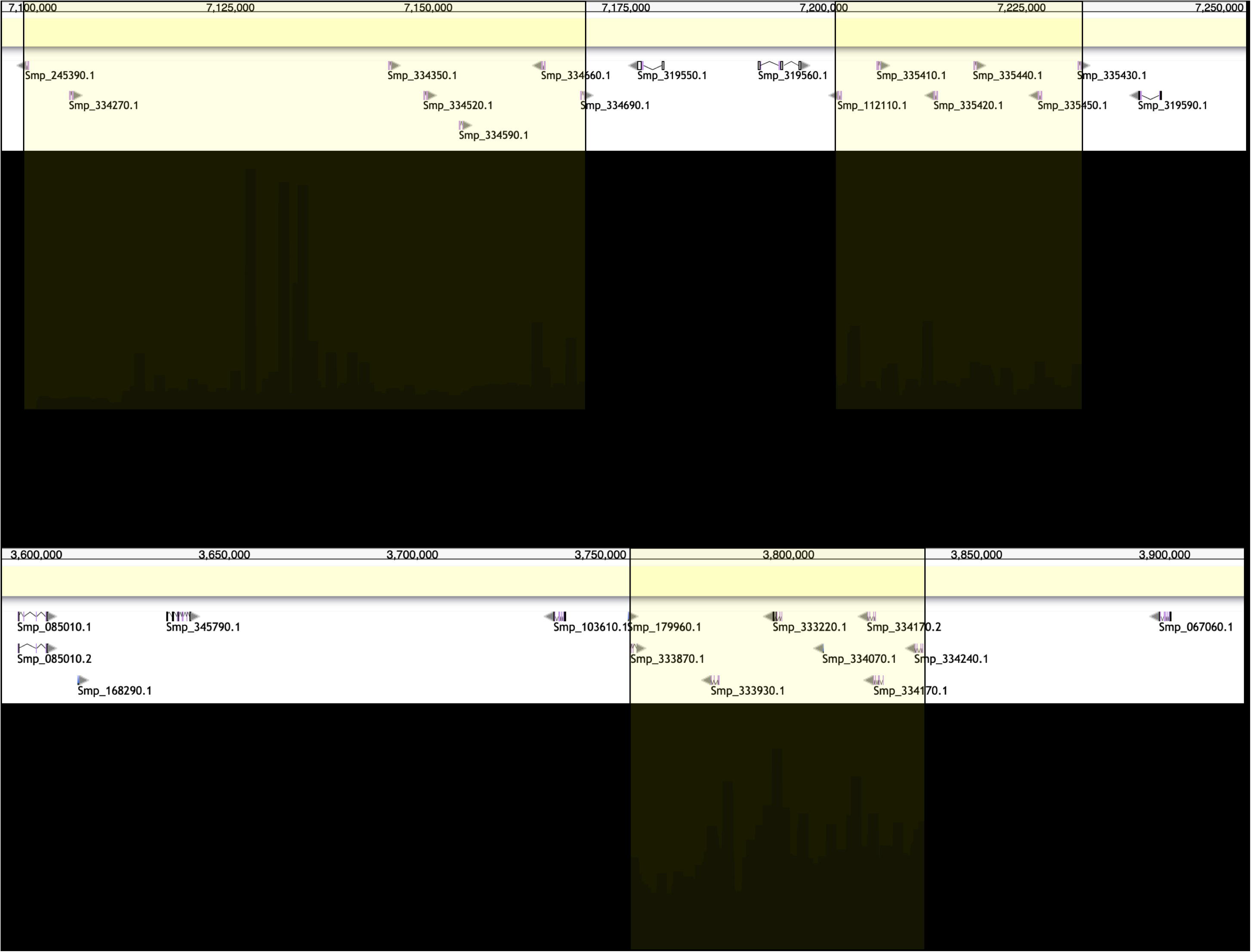
Resolving the repetitive IPSE and omega-1 loci. Genes in the (a) IPSE loci and (b) Omega-1 locus shown in v9 through gene model annotations (top tracks) and genomic coverage mapping (bottom tracks) with yellow boxes to connect gene annotations to genomic coverage. The annotations show the v9 gene models, some of which coincide with elevated read coverage. The histogram in the coverage plots show depth of read coverage and compared with the flanking regions, the depth is elevated in the IPSE and omega-1 loci suggesting these gene arrays are smaller in this assembly than their true size.

**Figure S3:**
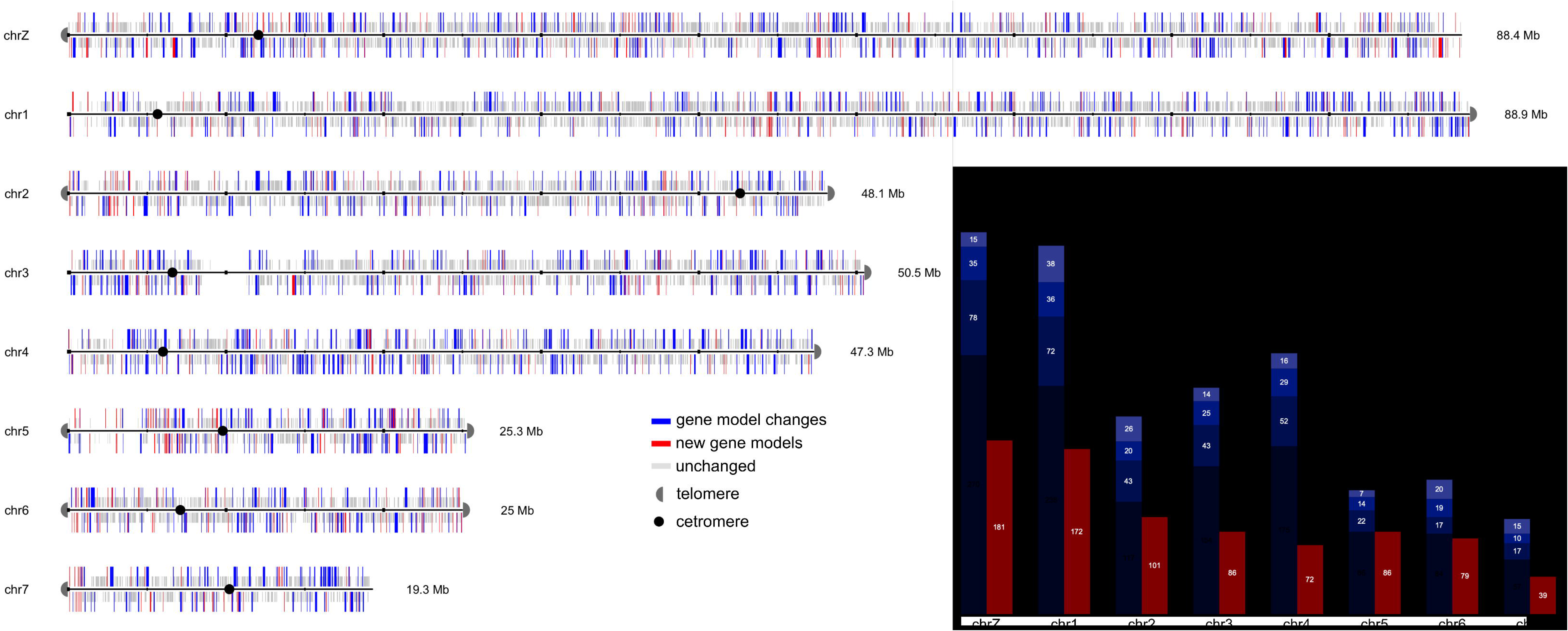
Gene changes from genome v5 to v9 of *S. mansoni*. There have been a total of 3,610 gene changes represented by 810 new, 867 deleted, 344 merged, 189 multiple copies, 190 split, and 1,210 structurally changed. The bar graph shows totals of different protein-coding gene changes in the primary assembly (i.e. no gene fragments, haplotypes, or pseudogenes).

**Figure S4:**
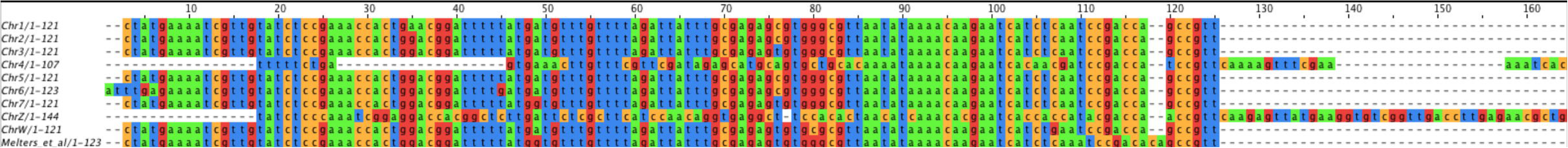
Alignment of the centromeric repeat sequences relatedness between all *S. mansoni* chromosomes. MAFFT/Jalview alignment of a single centromeric repeat unit from each chromosome shows high similarity between chromosomes 1-3, 5-7, and W. Chromosomes 1-3, 5-7, and W are 93.1-98.5% identical to a 123bp centromeric repeat proposed by Melters *et al* [26]. The centromeric repeats for chromosomes 4 and Z are diverged from the other chromosomes.

**Figure S5:**
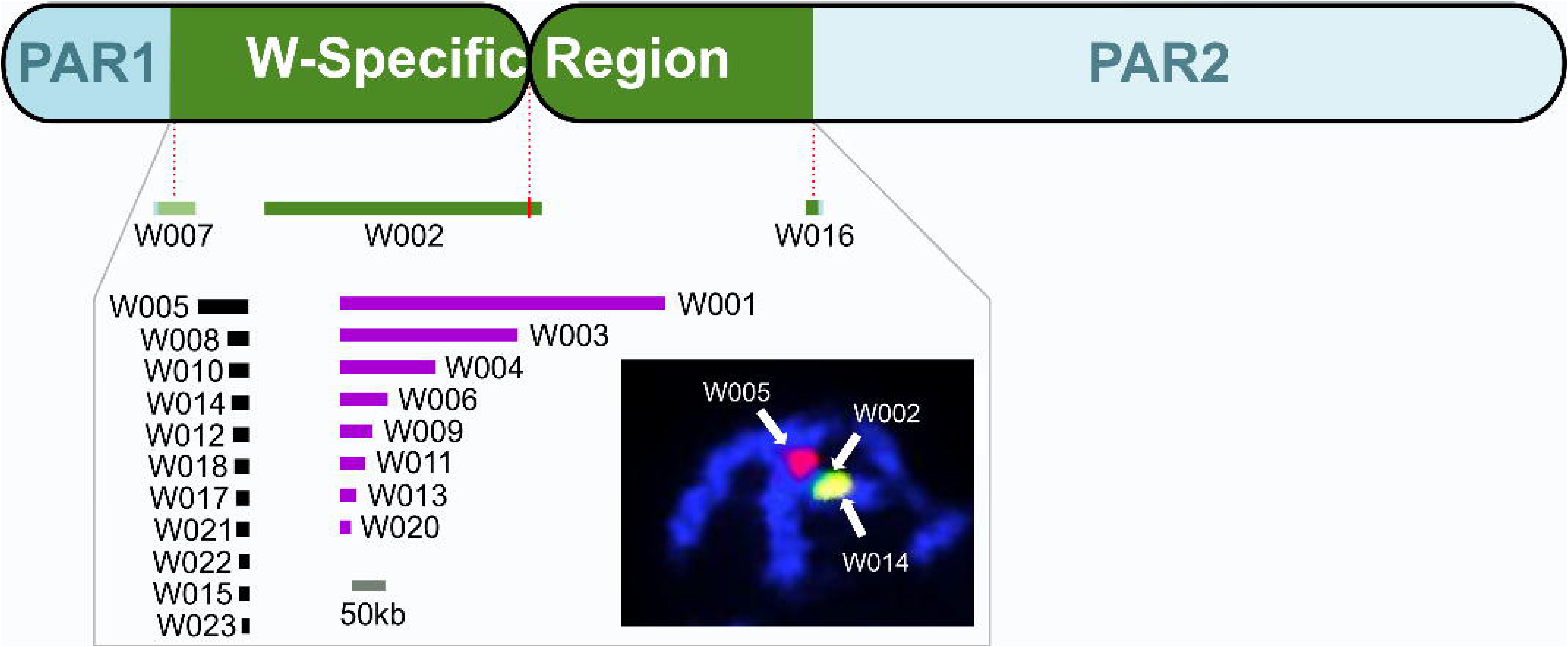
Validation of the assembly and placement of W-specific scaffolds using metaphase FISH. Twenty-two W-specific scaffolds existed after computational and manual assembly. Scaffold W007 contained the junction from PAR1 into the WSR and scaffold W016 spanned the WSR into PAR2. The centromeric repeat for the W chromosome was in scaffold W002 (7.65-7.75 Mb) with the orientation of this contig inferred from alignment of centromere sequence in this scaffold. The remaining 8 scaffolds with gametologues (purple) and 11 scaffolds without gametologues (black), whose positions and orientations could not be determined using sequence data alone, were placed using metaphase FISH.

**Figure S6:**
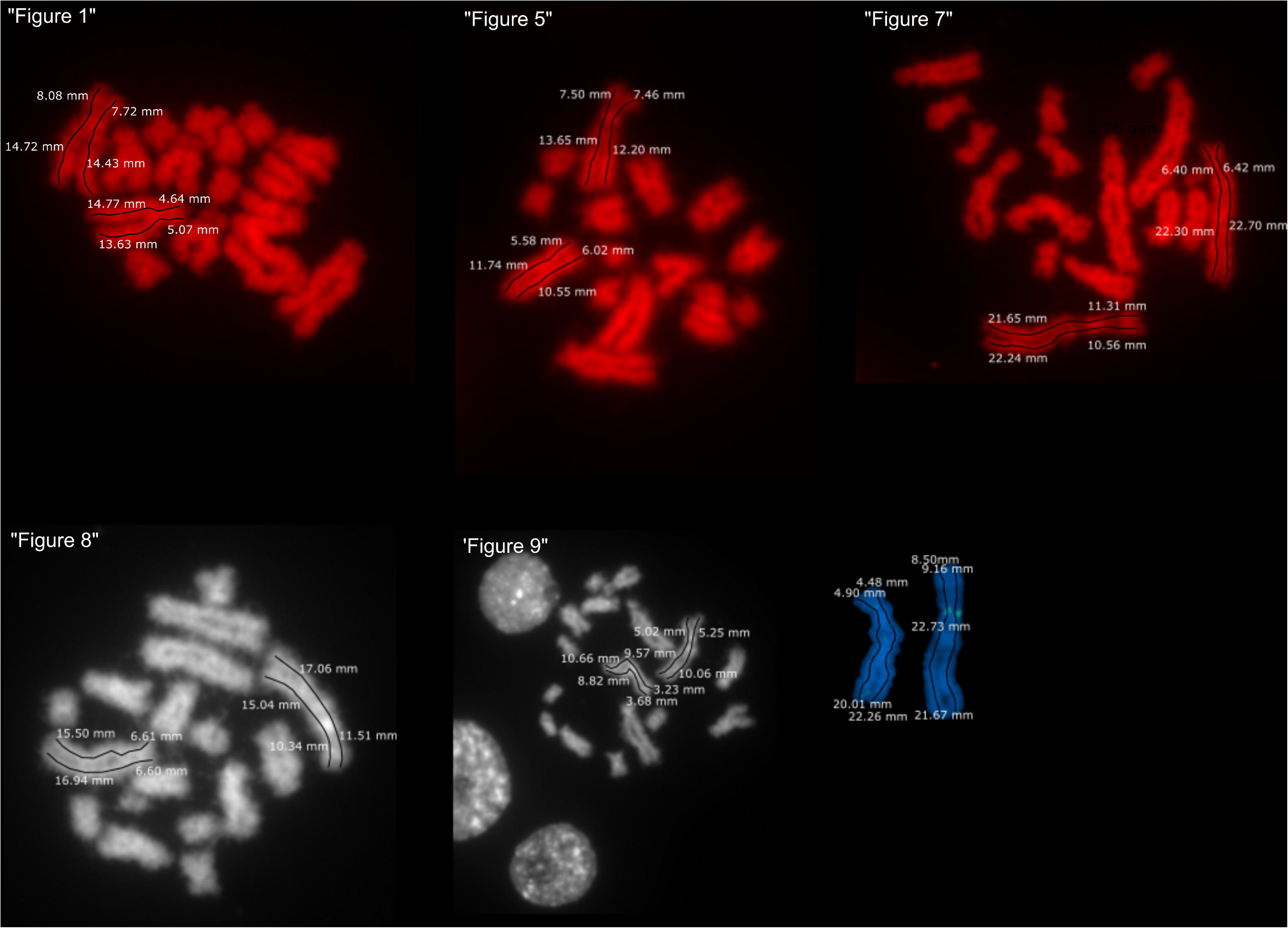
Measurements of Z and W chromosomes from 6 female metaphase cells. The W chromosome is approximately 14.7% larger than the Z chromosome based on measurements taken of the chromosomes from the metaphase figures shown. Measurements were taken using the measurement tool in Inkscape. This figure is consistent with previously published measurements from 22 female metaphase cells [33].

**Figure S7:**
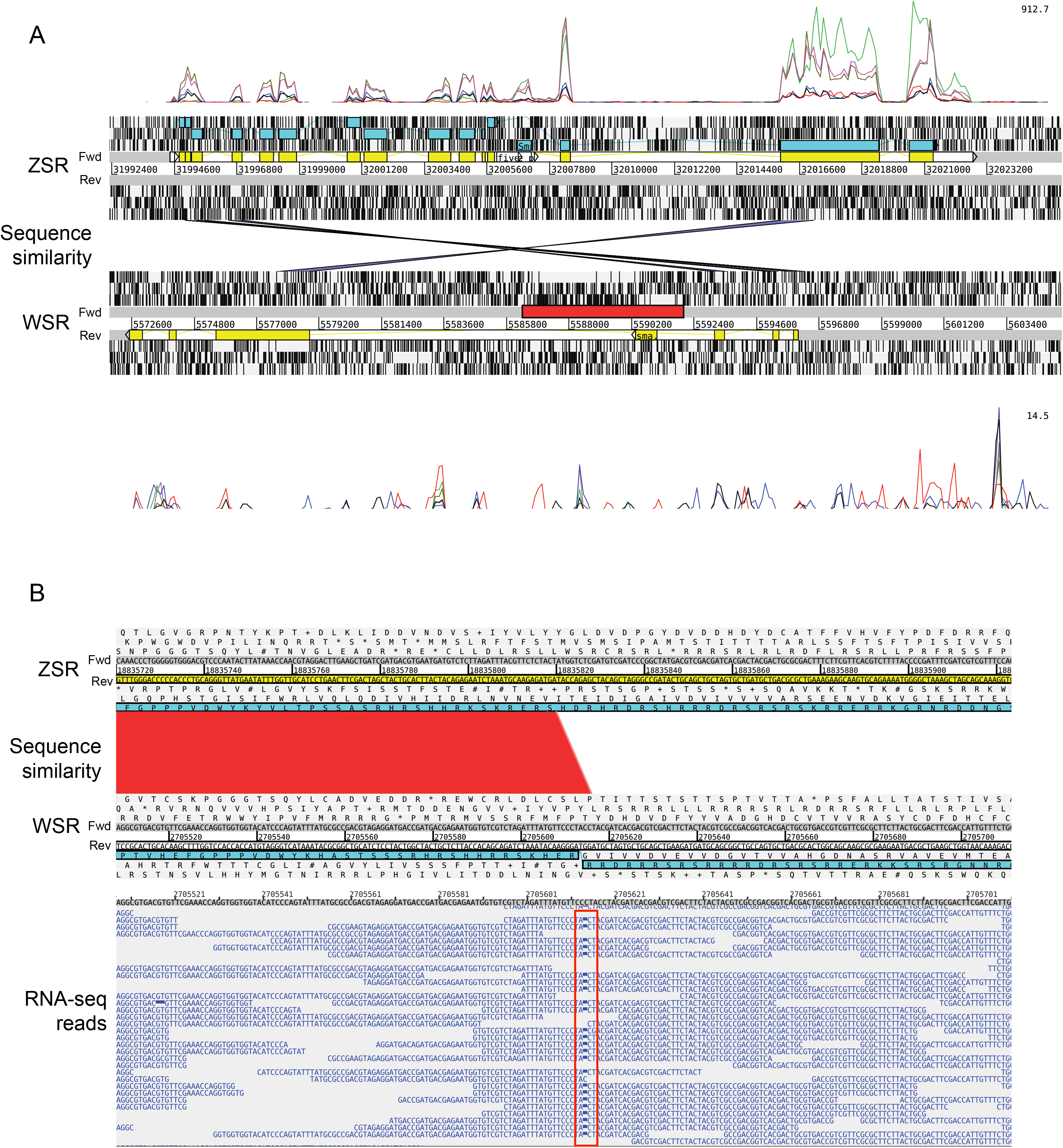
Comparisons of ancestral region gametologues between ZSR and WSR. (a) The Z gametologue Smp_158310 is clearly expressed in males (red RNA-seq coverage) and females (blue RNA-seq coverage), but the W gametologue Smp_318710 is not. Furthermore, the gene model is incomplete on WSR and there is a transposon inserted within the gene (red bar), resulting in a pseudogene. The genes are inverted between ZSR and WSR, indicated by the overlapping sequence similarity bars. (b) The genome sequence for the WSR gametologue of U2AF 65kDa (Smp_348790) subunit contains a single base insertion, suggesting a possible frameshift mutation. However, RNA-seq reads show that this is a sequencing error and the corrected gene model based on this data results in an N-terminal amino acid sequence more similar to, although still somewhat divergent from, the Z gametologue (Smp_019690).

## Notes

https://zenodo.org/record/5149023#.YRcKZtNKhQI

### Competing Interest Statement

The authors have declared no competing interest.

